# Aberrant newborn T cell and microbiota developmental trajectories predict respiratory compromise during infancy

**DOI:** 10.1101/736090

**Authors:** Alex Grier, Nathan Laniewski, Ann L. Gill, Haeja A. Kessler, Heidie Huyck, Elizabeth Carbonell, Jeanne Holden-Wiltse, Sanjukta Bandyopadhyay, Jennifer Carnahan, Andrew M. Dylag, David J. Topham, Ann R. Falsey, Mary T. Caserta, Gloria S. Pryhuber, Steven R. Gill, Andrew McDavid, Kristin M. Scheible

## Abstract

Neonatal immune-microbiota co-development is poorly understood, yet appropriate recognition of – and response to – pathogens and commensal microbiota is critical to health. In this longitudinal study of 148 pre- and 119 full-term infants from birth through one year of age we found that postmenstrual age, or weeks from conception, not post-natal age, is the dominant factor influencing T cell and mucosal microbiota development. Numerous features of the T cell and microbiota functional development remain unexplained, however, by either age metric and are instead shaped by discrete peri- and post-natal events. Most strikingly, we establish that prenatal antibiotics or infection disrupt the normal T cell population developmental trajectory, influencing subsequent respiratory microbial colonization and predicting respiratory morbidity. In this way, early exposures imprint the postnatal immune-microbiota axis and place an infant at significant risk for respiratory morbidity in early childhood.

**One Sentence Summary:** T cells and microbiota follow predictable, coordinated trajectories in newborns, and their coordination is an important determinant of illness in the first year.

**Graphical Abstract:** Nasal and rectal microbiome was assessed weekly in the NICU, monthly post-discharge, and when respiratory illness symptoms occurred. T cells were characterized at birth (cord blood), 36-42 weeks postmenstrual age, and at 12 months. Quarterly respiratory morbidity surveys determined the 12-month outcome of persistent respiratory disease. More frequent illnesses and chronic disease outcomes occurred by one year of age if development of these systems was discordant or disrupted.

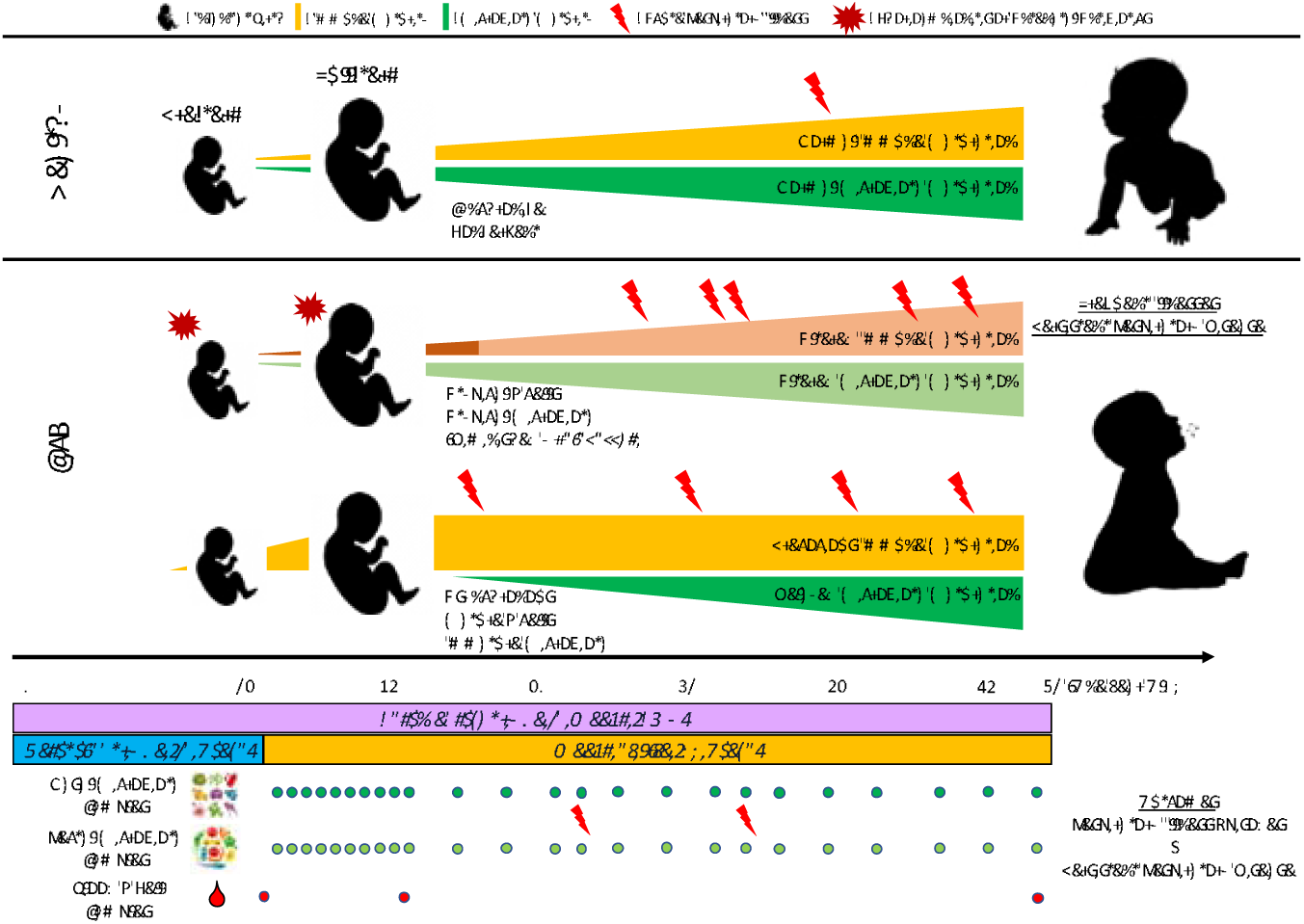

## Introduction

Function of the immune system and establishment of microbiota in human infants have a profound impact on subsequent health and disease. However, the factors influencing immune system and microbiota establishment and the extent of their inter-relatedness are incompletely understood (1-3). Normally, a developmental program determines major shifts in immune cell population maturation and distribution over the first 3 months of postnatal life (4). Though the exact stimuli for these shifts are unknown, it is increasingly clear that abnormal gut and respiratory microbiota during infancy associate with adverse outcomes such as atopy, stunted growth, and respiratory infection – outcomes that correspond to maladaptive immune system activity (5-9). Two recent studies demonstrated that the nasopharyngeal microbiome and virome together predict infant respiratory tract infection, but these cross-sectional studies left unresolved the sequence and inter-relatedness of the microbiome and immune development leading to such adverse events (10, 11). We propose that with health, there is a close integration of immune system development and microbiota establishment such that if either is perturbed by events like antenatal antibiotics, the outcome is poor health. The strength of the study presented here is that both systems are assessed together in a longitudinal fashion, considering multiple external factors and perturbations.

Many of the same extrinsic factors likely influence the parallel development of the immune system and the microbiome and it has become apparent that the systems have mutually formative effects on one another. The immune system, as it develops and as it’s maintained, is critically refined to determine what is self, what is commensal and what is pathogenic. Reported adverse effects following disrupted developmental processes support the concept of a critical neonatal window during which primary exposures and maladaptive immune responses risk lifelong health (12). Whether or not an exposure is differentially “remembered” by the immune system based on its timing during development, or if immune reprogramming alters a pattern of microbiota colonization, is not known, but such a concept is particularly relevant when considering the long-lived adaptive immune system. To date, there are no published longitudinal studies addressing how T cells and the mucosal microbiome are linked over time during early human development and none have explored the degree to which immunity and mucosal bacteria interplay impacts infant well-being in the first years of life. The need to understand the complex relationship between microbiota, the immune system and infant development is particularly urgent given the accelerating use of probiotic and prebiotic therapies in infants in the absence of adequate knowledge (13-16).

In this study, using a systems biology approach, postmenstrual age is identified as the dominant variable influencing patterns of T cell differentiation and gut and respiratory microbiota progression. However, several interdependencies between these systems could not be explained by their mutual dependence on age, but instead were traced to select perinatal exposures, and we observed significant relationships between aberrant microbiota-immune trajectories and respiratory illness in the first year. This work represents the first longitudinal assessment to date of the relationship between developing T cell populations and microbiota in human infants. Furthermore, it is the first to demonstrate a link between the co-development of these systems and clinical outcomes in infants.

## Results

### Study Design and Demographics

Neonatal subjects (n=267) born at 23-42 weeks gestational age (GA) were recruited within 7 days of birth at the University of Rochester from 2012-2016, as part of the NIAID-sponsored Prematurity, Respiratory, Immune Systems and Microbiomes study (PRISM) (Fig. 1A). In all, 122 preterm (PT, < 37 0/7 weeks gestation) and 80 full-term (FT, 37 0/7 weeks gestation) subjects completed the study to 12 months of age, corrected for premature birth, and were categorized as having or not having the primary outcome persistent respiratory disease (PRD) using previously published criteria (17). Cohort demographics are shown in Table 1. Sufficient blood to perform T cell phenotyping by flow cytometry was collected at three pre-defined timepoints, from 55% of subjects at birth (cord blood), 61% of subjects at NICU discharge and 38% at 12 months. For microbiota profiling, inpatient samples were obtained weekly and outpatient samples for PT and FT were obtained monthly, with additional sampling during acute respiratory illnesses. After sample processing, 16S rRNA sequencing, quality control, and removing subjects without immunophenotyping data, 149 subjects yielded 1748 usable nasal swab samples and 143 subjects yielded 1899 usable rectal swab samples. The median subject had 24 samples, with 28 days on average between samples. Finally, 109 and 117 subjects had sufficient combined T cell phenotyping and microbiota data to be included for immunome-nasal microbiota and immunome-rectal microbiota association analyses, respectively (Supplementary Tables 1-2).

**Table 1:** Subject Demographics

|  | Preterm (n=148) | Term (N=119) |
| --- | --- | --- |
| Gestational age (weeks) | 29.8 ± 3.7 | 39.6 ± 1.0 |
| Birthweight (g) | 1406.6 ± 620.8 | 3471.5 ± 511.4 |
| Female | 71.0 (48.0%) | 49.0 (41.2%) |
| Black or Asian race | 46.0 (31.1%) | 29.0 (24.4%) |
| Public Insurance | 81.0 (54.7%) | 59.0 (49.6%) |
| Maternal smoking postnatal | 29.0 (19.6%) | 15.0 (12.6%) |
| Delivered by cesarean section | 94.0 (63.5%) | 49.0 (41.2%) |
| Chorioamnionitis | 7.0 (4.7%) | 4.0 (3.4%) |
| Funisitis* | 35.0 (24.3%) | 0 (0.0%) |
| Pre-eclampsia | 26.0 (17.6%) | 0 |
| Antenatal steroids | 121.0 (81.8%) | 0 |
| Postnatal steroids | 47.0 (31.8%) | 10.0 (8.4%) |
| BPD | 25.0 (16.9%) | 0 |
| Supplemental O2 (Median FiO2 for first 14 days, IQR) | 21.9% (21-27.8) | 21% (room air only) |
| Postnatal infections (% with culture - positive bacteremia) | 24.0 (16.2%) | 0 |
| Received breastmilk (any) | 134.0 (90.5%) | 92.0 (77.3%) |
| Number of illness visits/subject (mean +/- SD) | 1.2 (1.9) | 1.0 (1.8) |
| Ventilator days (n) | 10.0 ± 18.3 | 0 |
| PRD** | 52.0 (42.6%) | 17.0 (21.1%) |

\*Funisitis calculated on 144 PT and 27 FT subjects (placental pathology available)
\*\*PRD measured in 122 PT and 80 FT

**Fig. 1.**
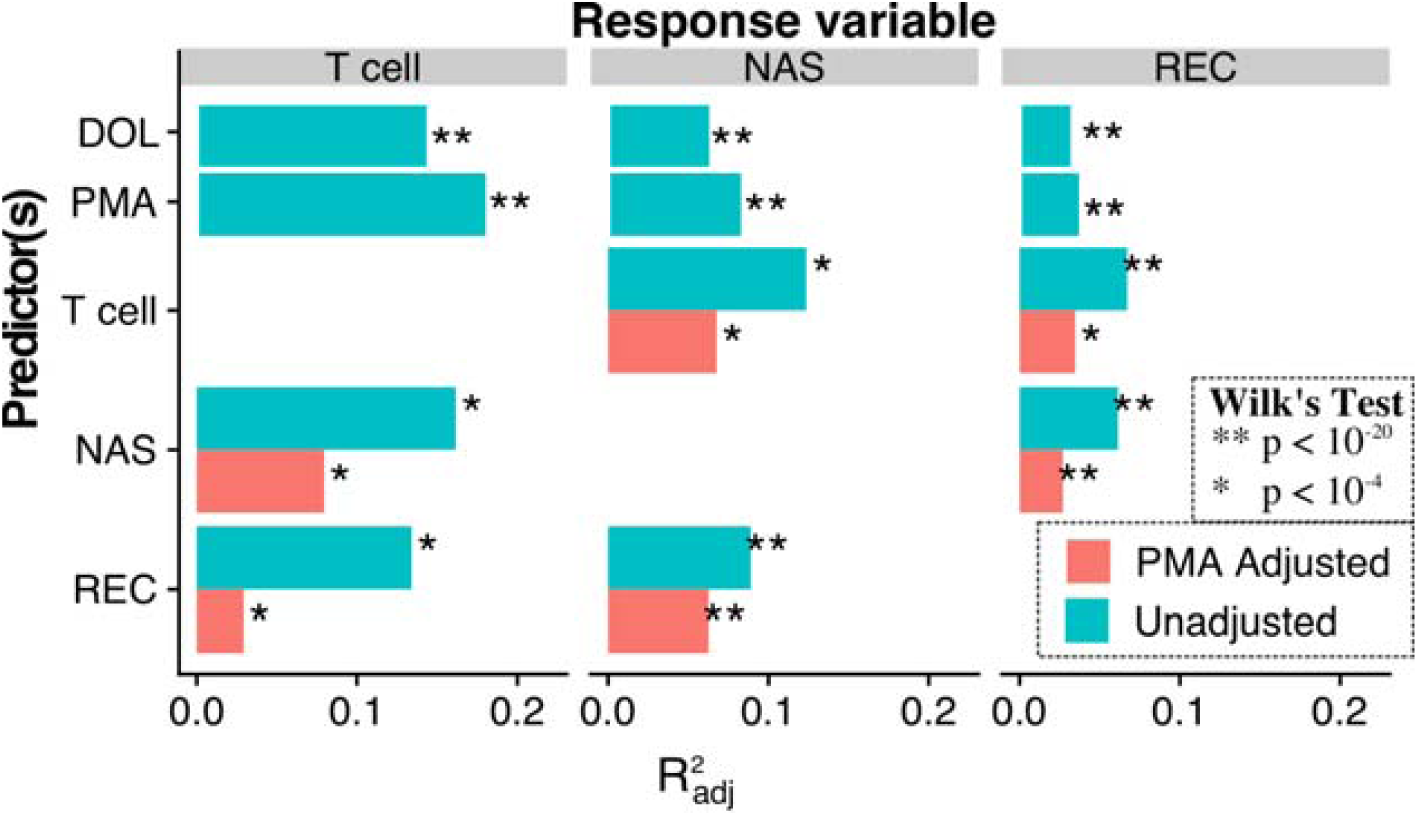
Systemic Interactions Between Age, T Cells, Microbiota and Age. The amount of variance in composition of nasal (NAS) and rectal (REC) microbiota and T cell immune populations that are explained by the predictors: postnatal day of life (DOL), postmenstrual age (PMA), T cell population composition and microbiota composition. Controlling for PMA, system interdependence was diminished but remained highly significant (red).

### Postmenstrual age exerts a greater influence on early T cell and microbiota development than does postnatal age

We predicted, based on our and other previous studies, that T cell and microbiota evolution in the first year of life would proceed in an age-dependent manner. Age can be defined in several useful ways in newborns, and these definitions have different implications in a cohort that includes infants born extremely premature. In considering the strengths of a combined PT and FT cohort, we tested two competing hypotheses regarding microbiota-immune maturation and infant age. First, that the microbiota and T cells would be shaped primarily by days of life (DOL), i.e., postnatal exposures would be the essential driver of T cell and microbial maturation. Under this hypothesis, we would expect PT and FT infants to display similar states at birth but distinct states if compared when both were at term equivalent postmenstrual age (PMA; defined as days since last known menstrual period, a proxy for time since conception), because the PT infants would be considerably older in DOL compared to FT newborns at that point. Alternatively, these systems could be more heavily influenced by an infant’s physiologic maturity, or PMA. If this alternative proved true, PT and FT subjects would exhibit distinct profiles at birth which would converge when PT infants achieved term equivalent PMA.

We applied unsupervised clustering approaches to reveal fine-grained, biologically interpretable categories of T cell populations and microbiota. The clustering algorithm FlowSOM identified 80 discrete populations of T cells using flow cytometry data (18): 50 from a T cell phenotyping panel (Tphe) and 30 from an intracellular cytokine panel (ICS). For the microbiota data, DADA2 was used to denoise and resolve the 16S rRNA amplicon sequence variants. We compared the effect of PMA and DOL on microbiota and T cell populations at a high level by using multivariate ANOVA to determine the explanatory power of each measure of age across all component microbes or T cell populations (Fig. 1). Compared to DOL, PMA was superior in predicting T cell, gut and respiratory microbiota composition (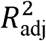 0.18, 0.04, and 0.08, respectively). Based on these results, we focused on PMA, rather than DOL, as the best predictor of T cell and microbiota maturation.

Anticipating that T cell populations and microbiota composition would show interrelated patterns of variation, we again applied multivariate ANOVA to quantify the amount of total variance the composition of one system could explain in another. All pairs of systems exhibited significant relationships with one another. The 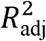 ranged from 0.05 (nasal microbiota explaining gut microbiota) to 0.15 (nasal microbiota explaining T cells). However, given that all systems exhibited strong associations with PMA, we reasoned that much of the observed effects would be due to the common influence of PMA progression within subjects rather than direct action of one system on the other. Indeed, adjusting for PMA in these models attenuated the variance explained between systems by approximately 50%, though all pairs were still significantly interrelated. These results support PMA as a significant driver of T cell and microbiota maturation, but further suggest a more complicated model in which these systems, albeit to a lesser degree, coordinate independently of host age.

### Premature birth transiently alters T cell development

Based on our finding that PMA (a variable that precedes birth and continues postnatally) better explains the temporal progression of T cell populations than DOL, we anticipated that PT and FT infants who were born at different PMA would begin with distinct phenotypes at birth but then converge as they achieved equivalent PMA. Using uniform manifold approximation and projection (UMAP) to visualize a reduced-dimension representation of each sample’s combined vector of ICS and Tphe T cell population proportions, we confirmed that PT and FT T cells clustered separately at birth, began to converge at 40 weeks PMA, and were fully overlapping between GA groups by 12 months (Fig. 2A, 2B).

**Fig. 2.**
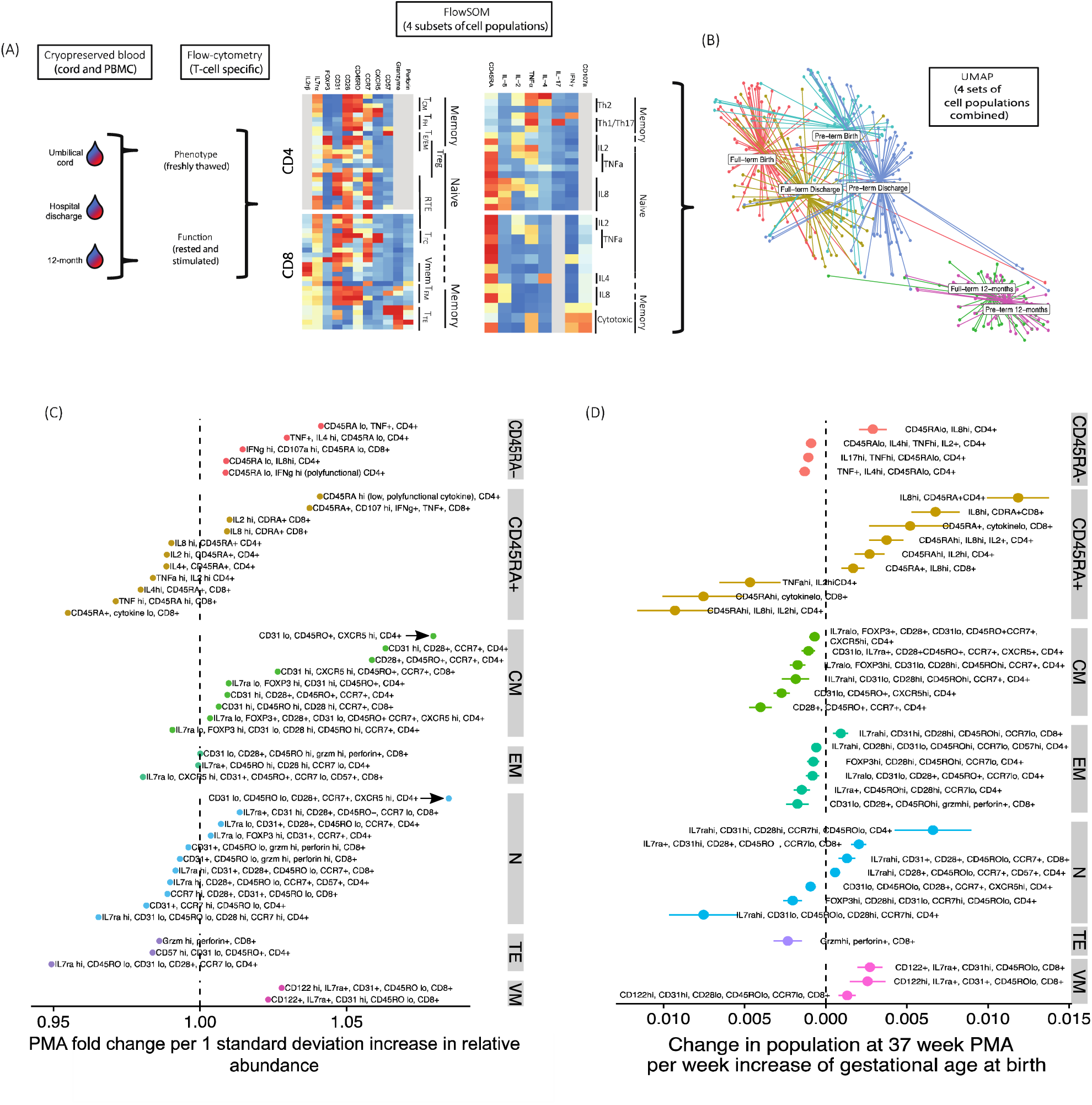
Early T Cell Development in Preterm and Full-term Infants Advances with Postmenstrual Age. (A) Flow cytometry was performed on blood obtained from 133 preterm and 79 full-term infants at birth, at the time of hospital discharge or at 36-42 weeks postmenstrual age (PMA), whichever occurred first, and at 12 months. T cells were characterized by phenotype (“Tphe”, unstimulated) and cytokine function (“ICS”, stimulated in vitro). Cell populations were identified using the Flowsom clustering algorithm. (B) Uniform manifold approximation and projection (UMAP) was used for dimensional reduction and summary visualization (each dot on the UMAP display represents a sample). Color overlays in UMAP show gestational age cohort and sampling timepoint. (C) Individual T cell populations that predict sample PMA are displayed, with populations that have inverse associations with PMA on the left of the dashed line, and positive associations to the right. The x-axis values indicate PMA fold-change per z-scored increase in the proportion of a subject’s cells assigned to that population. T cell phenotype subpopulations are grouped and colored based on CCR7 and CD45RO expression (TCM=central memory, TEM=effector memory, TN/SC-M=naive, stem cell memory, TTE= terminal effector) and CD122 (IL-2r□, VM=virtual memory). (D) T cell populations differentially abundant at 37 weeks PMA as a function of gestational age at birth are grouped by functional profile. The x-axis values shows the estimated change in population proportion at 37 weeks PMA, per one week change in gestational age at birth, and its 95% confidence interval, estimated by broken-stick regression models (see Methods). Populations are plotted left to right from those most abundant at 37 weeks in preterms to those most abundant in full-terms, with extrapolation occurring at GA > 37 weeks.

We next investigated the patterns of individual T cell population abundances over time from birth through one year. Previous studies focusing on a limited number of T cell subsets have suggested a straightforward model in which the fetal T cell developmental program progresses in a linear fashion from a naïve, tolerant state towards a more conventional memory and effector phenotype following repeat immune priming postnatally (19-22). To provide a more nuanced characterization of the PMA-directed T cell program, we examined populations that were selected as statistical predictors of PMA using elasticnet regression. Tphe populations were grouped according to the following established naming conventions: effector memory (EM, CD45RO+, CCR7-, CD28-), naïve (N, CD45RO-, CCR7+, CD28+), central memory (CM, CD45RO+, CCR7+), virtual memory (Vmem, CD8+, CD45ROlo, CD122hi), terminal effector (TE, CD45ROlo, CCR7-, CD28-), (23-28). ICS populations were first grouped into naïve and memory (CD45RA+ and CD45RA-, respectively), and then named based on predominant cytokine profile. Interestingly, T cells associated with the “youngest” PMA displayed TE marker combinations, including CD8+ T cells positive for cytotoxic granules. Naïve and EM subsets followed TE chronologically. Naïve populations showed considerable heterogeneity across PMA, indicating that a “naïve” T cell grouping by CCR7 and CD45RO expression is overly simplistic in characterizing early T cell development. CM populations were generally associated with older PMA, and several of the CM populations that arose earlier carried a FOXP3+IL7rαlow T-reg cell phenotype (Fig. 2C), again exposing the limitations in applying conventional T cell grouping using markers validated in a fully mature immune system. Functionally, CD4+ T cells progressed from naive TNF-α and IL-2 high, then to IL-8 high, then to polarized, polyfunctional cells at later PMA. Naïve, cytokine low and TNF-α positive CD8+ T cells were present at early PMA, then progressed through IL-4 and IL-8 positive, then cytotoxic CD45RA+. CD45RA low T cells were biased towards 12-month samples.

To isolate the impact of prematurity on T cell development, we performed a multivariate regression based on GA, PMA and interactions thereof, estimating GA-associated changes in abundance of T cell subsets at 37 weeks PMA (Figs. 2D and 2E). T cell subsets again were grouped based on CCR7, CD45RO, CD45RA, and CD122 expression as described above. Naïve subsets varied in their phenotype at 37 weeks across the GA range, and PT-associated naïve subsets were distinct in their lower CD31 expression and IL-8 expression. Memory and effector subsets (CD45RA-, CM, EM, TE) were associated with younger GA, with the exception of VM, which was associated with FT subjects (Fig. 2D). Even within the FOXP3+ populations, there was subtle variation in phenotype between PT and FT, with PT-associated Tregs displaying low CCR7 and CD31 expression.

We found an overall pattern of T cell convergence between PT and FT over one year, indicating that PMA plays a central role in early T cell maturation. However, some populations, such as the effector/memory T cells enriched in PT at birth, could instead be a transient response to perinatal inflammatory conditions. To explore this notion, we sought out cell population subsets with distinctly non-linear patterns of abundance from birth to discharge to one year. Utilizing the three timepoints typically captured for each subject, we identified 10 T cell populations with non-monotone V-or inverted V-shaped trajectories from birth to 12 months in PT samples (Supplementary Fig. 1). Most frequently, these population abundances followed a V-shaped trajectory that decreased sharply from birth, to 37 weeks PMA, followed by a slower recovery from 37 weeks to one year. This pattern was seen in several memory CD4+ and CD8+ T cell subsets, indicating a transiently activated T cell phenotype in PT at birth that resolves under more homeostatic conditions. Two CD4+ ICS populations, which were IL-8-positive, had inverted-V trajectories. Together these results reveal a trajectory in which pauci-functional memory, effectors and regulatory T cells are enriched during early development, and these give way to more “typical” naïve populations, followed by a gradual gain of fully functional memory T cell subsets.

### Atypical T Cell Developmental Trajectories Are Predicted by Inflammatory Exposures

As described, individual T cell population abundance was robustly associated with PMA, but individual populations are not easily interpreted in the context of a rapidly changing system. To characterize a more integrated, holistic T cell trajectory during infancy, we grouped Tphe and ICS samples into immune state types (ISTs) based on the abundances of their respective T cell populations using Dirichlet Multinomial Mixture (DMM) models (Fig 3). Each IST in this case represented an archetypal profile of T cell composition in terms of the abundance of the various T cell subpopulations, and samples were assigned to the IST which best explains their observed makeup. The seven T cell phenotype immune state types (Tphe ISTs, Fig 3A-C) and eight ICS immune state types (ICS ISTs, Fig 3D-F), numbered according to their average order of occurrence, exhibited strong associations with PMA (ANOVA, *R*^*2*^ = 0.86 and 0.69, respectively; Fig. 3C-3D). With the exception of one 12-month sample in ICS4, ISTs Tphe1-Tphe4 and ICS1-ICS4 were only seen in samples drawn at birth and discharge.

**Fig. 3.**
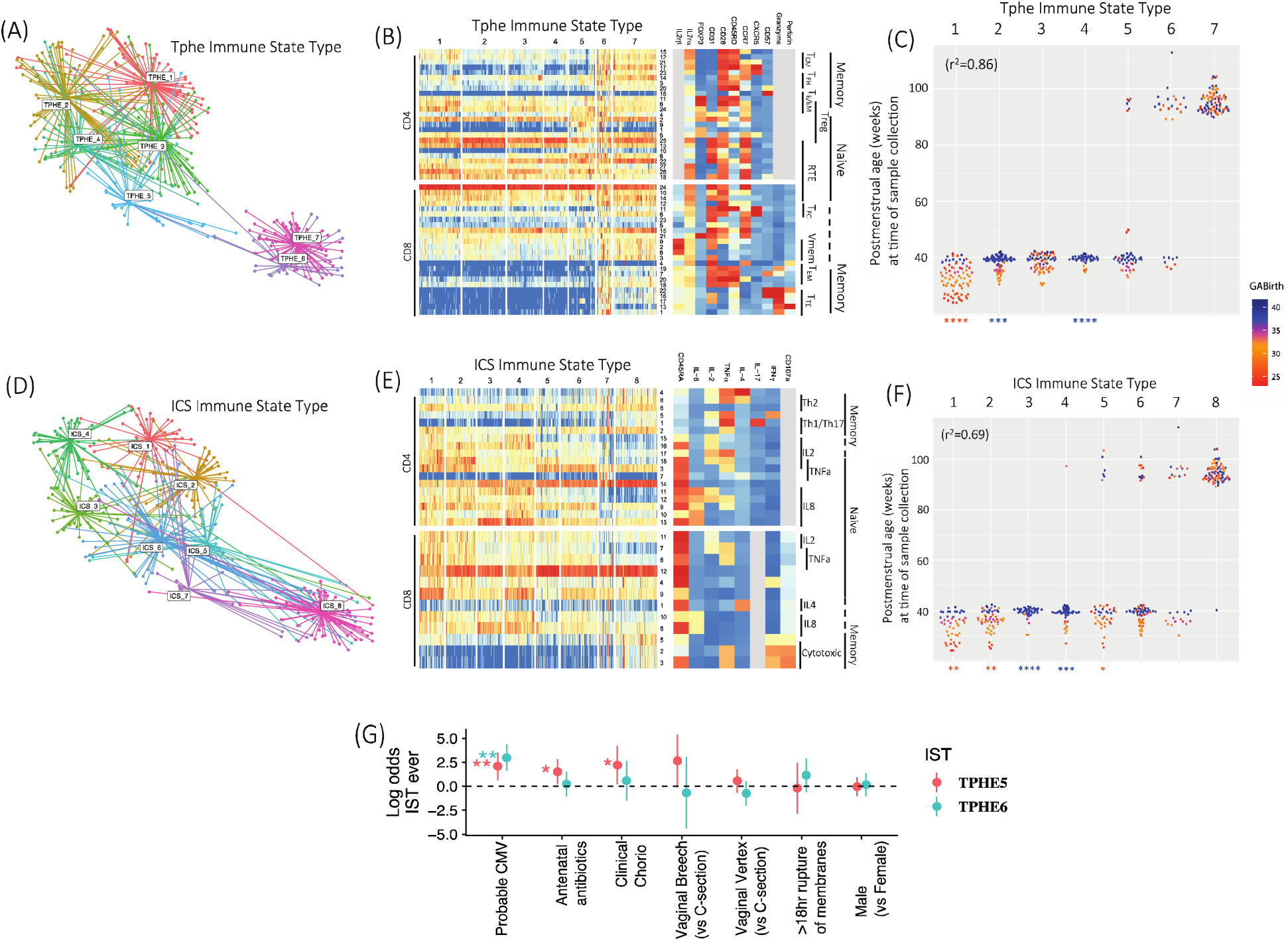
Immune State Types (ISTs) Advance with Postmenstrual Age, and Are Perturbed by Specific Clinical Exposures. T cell phenotype (A, B) and T cell function (D, E) immune state types (ISTs) were defined based on T cell population abundances and used to color UMAP projections of all samples. ISTs were enumerated according to the average postmenstrual age (PMA) at with they occur. Colors reflect relative abundances of component cell populations (rows); functional annotations and defining markers are shown in the heatmaps on the right. (C) Tphe and (D) ICS samples are plotted in the column corresponding to their IST, with PMA at sampling along the y-axis, and colored by gestational age at birth (GA). R2 values show correlations between IST and PMA. Asterisks at the base of the dot plots indicate significant enrichment for either preterm (orange) or full-term (blue) samples within an IST. (G) Joint logistic regression showing the log odds of ever being in Tphe5 or Tphe6 given exposure indicated, controlling for gestational age (*p<0.05, **p<0.01, ***p<0.001, ****p<0.0001).

Two Tphe ISTs (Tphe5 and Tphe6) deviated from the normal IST progression, and we hypothesized that their occurrence was associated with clinical exposures common to PT and FT, as opposed to development alone. In support of this, we noted that Tphe5 had marked Treg heterogeneity (variable CD31+, CD45RO-, CCR7-, CD28-, effector-like subpopulations). Several inflammatory or infectious conditions, including chorioamnionitis and exposure to antenatal antibiotics raised the odds of a subject ever entering Tphe5 by 9-fold (95% CI 1.2-66, p<.04), 4.5-fold (95% CI 1.3-16, p<.02) respectively, in a joint logistic regression model that adjusted for GA, sex, race, mode of delivery, and premature rupture of membranes (Fig. 3E). High abundance of CD57+ and cytotoxic CD8+ T cells in Tphe5 and Tphe6 raised our suspicion for cytomegalovirus (CMV), a known driver of T cell exhaustion and high CD57 expression (29, 30). Evidence for CMV infection was detected, by either serology or PCR at 12 months in 18 subjects overall, and in less than 8% of Tphe6 or Tphe5 negative subjects. No subjects had stigmata or evidence for congenital CMV infection. However, 40% of subjects ever entering Tphe6 (20-fold odds, p<.0001) and 19% of subjects ever entering Tphe5 (8-fold odds, p<0.005) tested positive for CMV. Tphe6 did not associate significantly with other variables tested. When restricting attention to subjects with Tphe6 at 12 months, the CMV associated persisted, but was attenuated and no longer significant when considering only subjects in Tphe6 at birth or discharge, suggesting that CMV exposure may cause Tphe6. In contrast, the Tphe5-CMV association remained significant even among subjects in Tphe5 at birth and discharge, despite no confirmed cases of congenital CMV. Indeed, having Tphe5 early in life increased by 5-fold (p<.001) the odds of having Tphe6 at 12-months, but having Tphe6 early in life exhibited no association with Tphe5 status. It is tempting to conclude in this case that Tphe5 itself is a risk factor, or a proxy for risk factors that precede CMV infection. ICS ISTs did not exhibit significant associations with any variables considered. Through modeling the normal T cell phenotype trajectory during infancy as a function of PMA, aberrant trajectories are revealed that indicate an infant’s prior exposure to and/or future risk to inflammatory or infectious conditions. On the other hand, T cell functional maturation, as measured by gain in polarized and polyfunctional cytokine profile, may be less impacted by early, *nonspecific* exogenous inflammatory exposures.

### Convergent Microbiota Community Progression Parallels T Cell Development In Preterms and Full-Terms

Having characterized the compositional progression of T cell population profiles with respect to PMA, we performed a similar assessment of the microbiota. To summarize and examine the data structure broadly, unweighted Unifrac distances between all samples within each body site were computed as a measure of β-diversity and were used to perform principal coordinate analysis (PCoA). For both body sites, the first principal coordinate (PC1) corresponded to PMA (Supplementary Fig. 2). Samples lower in PC1 tended to be taken prior to 40 weeks PMA, while advancement along the PC1 axis corresponded to an increasingly balanced proportion of PT and FT subjects. These results establish broad parallels between the developmental pattern of T cell populations and microbiota with respect to PMA.

To summarize microbiota composition and facilitate subsequent comparative analyses, we applied a similar approach as performed on T cell populations using DMM modeling to partition samples into characteristic community state types (CSTs) based on their compositional profiles. Based on model fit and parsimony, 13 CSTs were defined for both respiratory (nCST) and gut microbiota (gCST) and were enumerated (1-13) according to the average PMA at which samples assigned to each CST were collected. Both gCST and nCST 1 were predominated by *Staphylococcus*, which was replaced over time with more niche-specific taxa in later CSTs, including *Enterobacteriales* and *Clostridiales* in the gut and *Streptococcus* and *Corynebacterium* in the respiratory tract (Fig. 4A-4B). Progression from CST 1 to 13 in the gut and the nose was strongly associated with PMA (ANOVA, *R*^*2*^ = 0.57 and 0.61, respectively) (Fig. 4C-4D). Several early CSTs with the lowest average PMA were dramatically over-represented by PTs, again suggesting a unique PT microbiota. Overall though, PT and FT subjects tended to converge to a shared microbiota and most CSTs were represented by equal proportions of FT and PT infants. However, a small number of gCSTs and nCSTs occurring later in the first year of life violated this tendency and were overrepresented by either PT or FT subjects. For example, gCST 9 contained 78% PT samples vs. 22% FT samples (p<0.05, two-tailed binomial test) and was notable for diminished levels of *Bifidobacterium* relative to gCSTs 8 and 10, which occurred over similar PMA intervals but which were over-represented by FT subjects (p<0.01 and p<0.05, respectively). These findings reveal that while the sequence of CST occurrence depends primarily on PMA, maturity at birth biases infants towards or against entering certain CSTs, even months after birth.

**Fig. 4.**
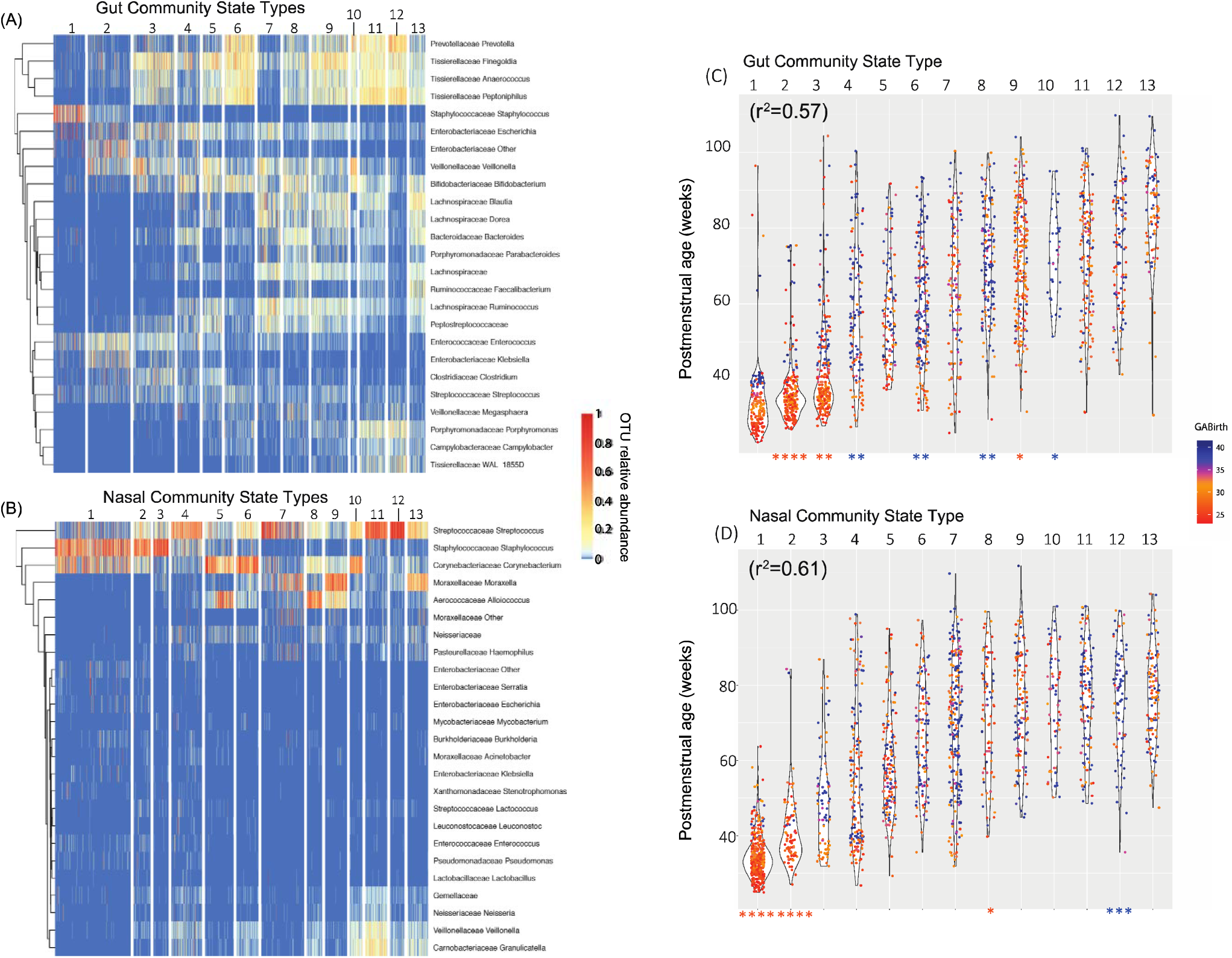
Premature Birth Influences Long-Term Age-Related Respiratory and Gut Microbiota Community Progression. Microbiota community profiling was performed on (A, C) rectal and (B, D) nasal samples obtained from 159 infants during regular surveillance and acute respiratory illness. (A, B) Microbiota community state types (CSTs) were defined for each body site based on sample composition, and the relative abundances of the top 25 most abundant genera were visualized using heatmaps, with samples as columns, clustered by CST. CSTs were numbered according to average PMA of occurrence. (C, D) Samples within each CST were plotted against subjects’ PMA at the time of sample collection. Each dot represents a single sample, colored by the subject’s GA. R^2^ values show correlations between CST and PMA. Asterisks at the base of the dot plots indicate significant enrichment for either preterm (orange) or full-term samples (blue) within a CST (*p<0.05, **p<0.01, ***p<0.001, ****p<0.0001).

### T Cells and Microbiota Interact Sparsely After Controlling for PMA

The strong relationship between PMA, T cell and microbiota state type trajectories suggests that the immune system and microbiota are regulated in tandem with the developing infant mucosal ecosystem. We therefore asked if T cell-microbial interactions occur beyond what can be explained by host age and whether such interactions might imperil an infant’s health. In order to explore this question, we modeled overall CST *duration, occurrence ever*, and *time to first occurrence* as functions of T cell population abundances and state types at specific time points. In the *duration* model, the number of days a subject spent in a given CST, adjusting for the total length of surveillance, was the response. For the *occurrence ever* model, the log odds that the CST ever occurred in a subject was the response. Lastly, in the *time to first occurrence* model, the response was the DOL of the subject’s initial transition into the CST. We modeled each as a function of one of the immunological parameters (IST or T cell population abundance at a particular time point), and adjusted for GA and mode of delivery. This approach reduced the longitudinal time series down to a sequence of subject-level summaries amenable to typical cross-sectional analyses. We fit models on all pairwise combinations of CSTs and immunological parameters. The significant results of these tests, which corresponded to interactions between the immune system and microbiota present in our cohort, were visualized as networks (Fig. 5A, Supplementary Fig. 3). Among the models of CST duration, of the potential 6318 possible associations between the 26 CSTs and 243 immunological parameters, only 10 Tphe and no ICS cell populations achieved statistical significance after multiple test correction. CST-associated CD4+ populations preceded, but CD8+ populations followed, the average onset of their associated CST, suggesting a temporal directionality to these relationships. For the models of CST occurrence ever, of the 15 total ISTs, only 3 (Tphe1, Tphe3 and Tphe5) were significantly correlated with a single CST (nCST 8). The most striking finding in the network was that entry into Tphe5 by the time of hospital discharge (n=25 subject-samples) precluded a subject ever entering into nCST 8, which itself was only ever observed after hospital discharge (Fig. 5B).

**Fig. 5.**
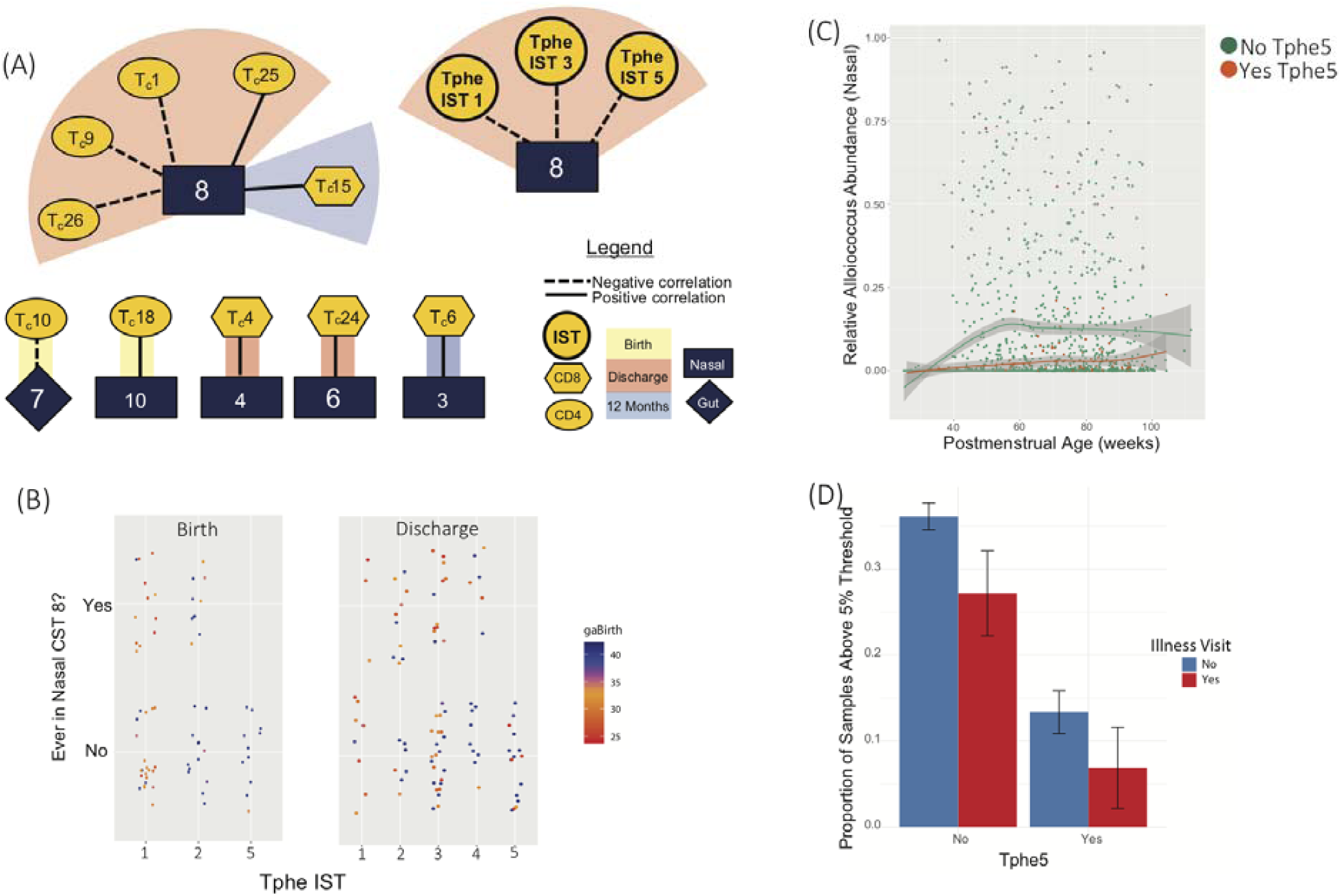
An early maladaptive immune state type precludes *Alloiococcus* colonization, which increases disease risk. (A) Time spent in microbial community state types (CSTs) was modeled as a function of T cell features (using both individual T cell population abundances and immune state types (ISTs) at each time point), controlling for gestational age (GA) at birth and mode of delivery, to test for significant microbiota-immune associations. Blue squares/diamonds show nasal/gut CSTs and yellow circles/hexagons/ovals show ISTs/CD8 populations/CD4 populations respectively. Connected shapes indicate significant association at 10% FDR. Background color indicates the timepoint at which the immune feature was associated with the connected CST. (B) Subjects whose samples were categorized as Tphe5 at birth or discharge time points never subsequently entered nCST 8. Sample color indicates gestational age at birth. (C) Relative abundance in the nose of nCST 8 dominant genus *Alloiococcus* (y-axis) vs.PMA of sampling (x-axis), is shown for subjects with (orange) or without (green) Tphe5 IST at birth or discharge. (D) Bar height indicates the proportion of nasal samples (+/-95% C.I.) with *Alloiococcus* relative abundance > 5% grouped by the presence of Tphe5 at birth or discharge, further segregated by sampling at illness (red) and surveillance (blue) visits.

In the time to first occurrence model, a greater number of gCSTs, CD8+ T cell populations and ICS cell populations exhibited significant associations than in the duration and occurrence ever models (Supplementary Fig. 3B). Infants who were delayed in their entry into the *Streptococcus*-dominant nCST 4 had higher TNF□or IFN□+ naïve CD8+ T cell frequencies at discharge and one year. Higher frequencies of effector CD8+ populations were also found in subjects with delayed gCST 9 entry (*Bifidobacterium* and *Bacteroides* low). Alternatively, the occurrence of gCST 3 was accelerated in subjects exhibiting Tphe2 at discharge (Supplementary Fig. 4). gCST 3 was the most diverse and mature gCST prior to discharge, and the earliest gCST in which *Clostridia* are prevalent. Notably, our previous study shows that the early presence of *Clostridia* in newborns predicts better growth velocity(6). Overall, the sparsity of these associations underscores the predominant role that host age plays in driving the abundance of T cell populations and microbiota composition. However, significant relationships between T cells and microbiota do occasionally occur even after accounting for the influence of host age, and the sequence of occurrence of associated immune markers and microbial CSTs relative to one another imply bidirectional imprinting between these two systems.

### Increased Risk for Respiratory Morbidity Following Disruption of Immune-Microbiota Axis by Antenatal Inflammatory Exposures

Closer examination of the infrequent but significant T cell-microbiota associations revealed the involvement of a bacterial genus with previously described clinically relevant functions. nCST 8, which was common overall but never observed in infants who manifested the Tphe5 immune state at either the birth or discharge sampling timepoints, was dominated by *Alloiococcus. Alloiococcus* was virtually absent in samples collected prior to hospital discharge, but appeared soon after, maintaining stable mean abundance through one year. Previous reports indicate that in children, *Alloiococcus* in the respiratory tract is negatively associated with acute respiratory infections (31, 32). Our results show that *Alloiococcus* is the distinguishing feature of nCST 8, and the occurrence of nCST 8 was precluded by the presence of antenatal inflammation-associated Tphe5 in early life. We therefore sought to assess the relationship between *Alloiococcus* abundance in the nose, acute respiratory illness, and early immunophenotype, controlling for multiple confounders. To identify episodes of respiratory illness post-NICU discharge, infants were scored by parents using a self-reported modified COAST score when respiratory symptoms arose (33).

As expected, the Tphe5 immunophenotype at birth or discharge was associated with diminished *Alloiococcus* abundance in the nose across all post-discharge timepoints, yielding a 7-fold reduction (3-14 fold, 95% CI, p-value < 0.001; Fig. 5C), while controlling for DOL, GA, mode of delivery, and repeated sampling of subjects. Additionally, for every 10% increase in *Alloiococcus* relative abundance within a sample, there was a 1.4-fold reduction in the odds of the sample having been taken during acute illness compared to healthy surveillance (1.1-1.8 fold, 95% CI, p-value < 0.003), controlling for confounders as above. Considering the joint effects of acute illness and Tphe5 occurrence at birth or discharge as predictors in the same model, we found that both were associated with reduced *Alloiococcus* abundance, (log ratios −0.9 ± 0.4 and −1.9 ± 0.8, respectively, 95% CI; p-values < 0.001; Fig. 5D). However, despite negative associations between Tphe5 and *Alloiococcus* abundance, and *Alloiococcus* abundance with illness, Tphe5 was not significant as a predictor of illness, either by itself (log odds = 0.5 ± 0.7, 95% CI, p-value = 0.13) or in conjunction with *Alloiococcus* relative abundance (log odds = −0.4 ± 0.7, 95% CI, p-value = 0.33), controlling for confounders in both cases. Taken together, these results show that bidirectional T cell-microbiota relationships occur infrequently, but when present, can be strongly linked with critical health outcomes. Furthermore, the temporal progression from prenatal inflammatory exposure to T cell phenotype to microbiota to clinical outcome suggests that the cascade of events leading to disease states in infancy is initiated early and involves a complex but observable interplay between exposure, host response and microbial development.

### Atypical Immune and Microbial Developmental Trajectories Predict Respiratory Morbidity

Observing that rare T cell-microbiota interactions occurring independently of PMA impacted respiratory morbidity led us to hypothesize that mistimings in development of T cells or microbiota increased the risk of chronic respiratory morbidity in the form of PRD. To test this hypothesis, we developed a quantitative model of the “normal” relationship between PMA and T cell and microbiota composition. We trained two sparse regression models that used the T cell populations and microbiota abundance vectors to predict log2-transformed PMA at sample collection. Holding out a subject’s longitudinal record, the cross-validated models strongly predicted PMA using either T cell populations (R^2^=0.77) or bacterial taxa (R^2^=0.65) (Fig. 6A). For each subject, the fitted intercepts of these models, which here represent the predicted PMA at 37 weeks actual PMA, indicate the subject’s microbiota and T cell maturity relative to “normal” at 37 weeks PMA (see “*Immunological and microbial developmental indices*” Methods for details). The fitted slopes of the models indicate a subject’s *rate* of microbiota and T cell maturation over the first year, again relative to normal. These four fitted parameters define a *developmental index* (DI) for each subject, which was used to assess mistiming with respect to age, or asynchrony between age, T cell and microbiota development.

**Fig. 6.**
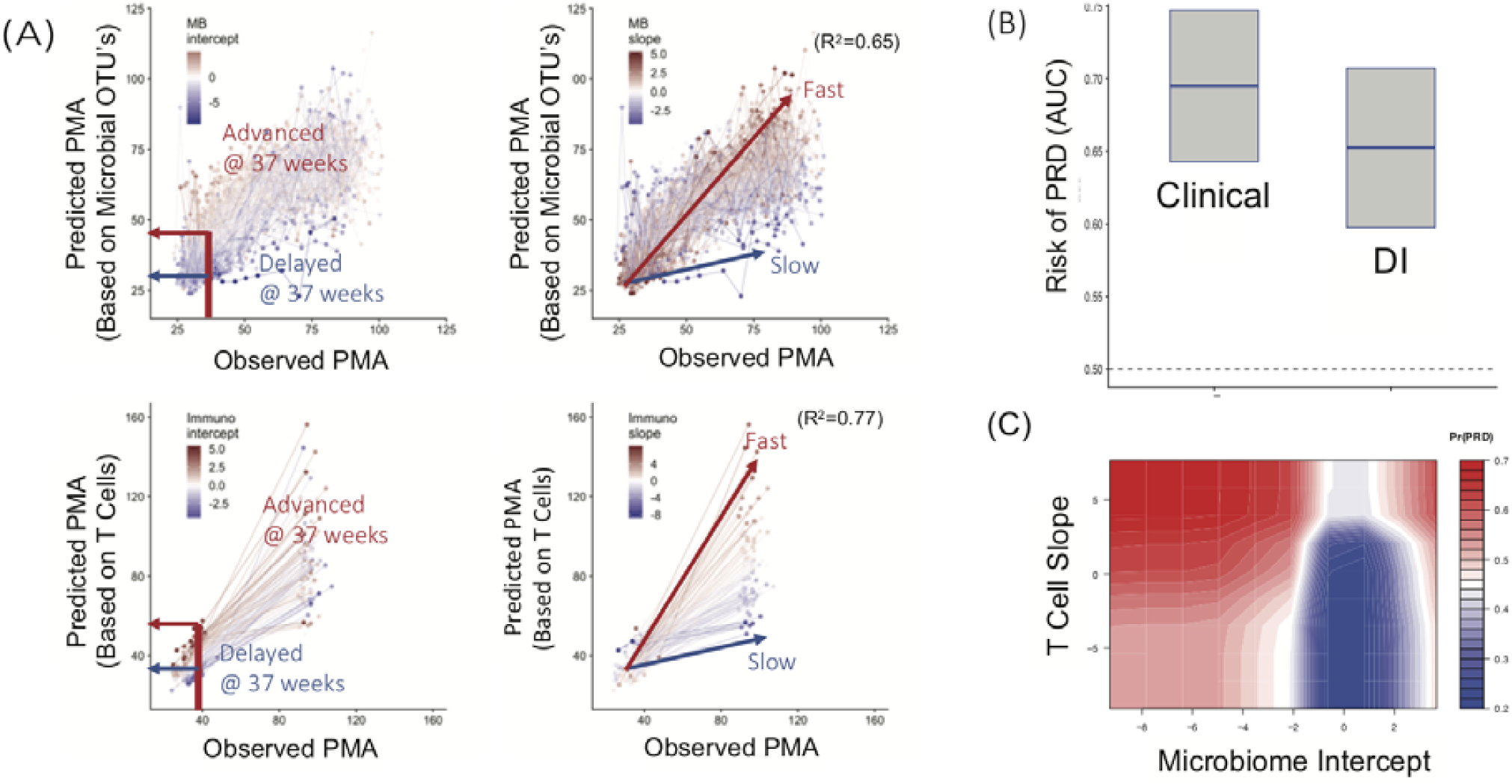
Mistimed Immune and Microbial Development Predict Respiratory Outcome. Elastic net regression predicts a postmenstrual age (pPMA) based on both T cell populations and microbial operational taxonomic units (OTUs), separately. (A) The pPMA of a subject is plotted against the observed age (oPMA) at the time of sampling to establish an intercept at 37 weeks (left) and a slope (right), corresponding to maturity at term equivalent and rate of maturation over the first year, for both T cell and microbiota. Z-scores for each subject’s slope and intercept are indicated as color overlays (red - relatively advanced maturation at term equivalent/faster development, and blue - relatively delayed maturation at term equivalent/slower development). A “Developmental Index” (DI) was constructed using these four parameters: the z-scores of T cell and microbiota intercepts and slopes. (B) A random forest machine learning algorithm predicts persistent respiratory disease (PRD) from known risk-factor clinical variables and from the T cell and microbiota-based DI. Box plots show the area under the curve calculated for each set of variables (mean and CI). (C) The contour graph demonstrates the two DI components, the microbiota intercept and T cell slope, with the best predictive strength for PRD risk, controlling for clinical factors. Blue = lower PRD risk, red = higher risk.

We used random forest classification models to compare the predictive power of the DI alone to that of a set of known clinical risk factors for PRD. The clinical features were race, maternal education, sex, GA, birthweight, season at birth and oxygen supplementation integrated over the first 14 days of life (34). The four DI features were the z-scores of the microbiota and T cell slopes and intercepts. In cross-validation, the clinical features predicted PRD with area under the curve (AUC) of 0.69 (0.59-0.79 95% CI). The features contributing most to the outcome were increased oxygen exposure, lower birthweight and younger GA (Supplementary Fig. 5).

Notably, when compared to clinical predictors, the developmental index had statistically equivalent skill in predicting PRD (Fig. 6B, AUC 0.64, 0.54-0.74 95% CI). Combining the clinical features and the developmental variables did not improve the AUC of the predictive model, further evidencing that T cell and microbial development may have durable effects on health outcomes that are equal in their impact to traditional clinical characteristics. Of the four components generating the DI, the microbiome intercept and immune slope had the largest variable importance scores. In exploring the functional relationship between PRD and these factors, we observed that an immature microbiota at term equivalent PMA increased the risk of PRD by over 2-fold, and this effect was magnified in subjects with accelerated T cell maturation (Fig. 6C). These results indicate that the timing of T cell and microbiota maturation *relative to an infant’s age* plays an integral role in promoting or undermining respiratory health.

## Discussion

Birth marks the commencement of a dynamic interplay between innate developmental programming, colonization and assembly of the microbiome, and differentiation and maturation of the adaptive immune system which influences health from infancy through adulthood. In healthy infants, this process balances the accommodation of commensal microbiota, appropriate immune response to pathogens, and functional maturation of the organs at the mucosal interface between human host and environment. Previous studies have generally applied a cross-sectional approach to summarizing microbiota and immune systems in infants, which precludes the ability to probe potential causal factors or clinical consequences of the correlations. By creating new longitudinal models of microbiota composition and T cell populations, we were able to establish conceptually and analytically tractable representations of the co-development of these systems, and in so doing, revealed several key findings. First, T cells and microbiota exhibit structured patterns of progression synchronized by PMA, with pronounced differences between PT and FT infants in very early life, but a tendency towards convergence postnatally. Furthermore, outside the framework of development driven by PMA, interactions occur between T cell population profiles and microbiota community structure, with early atypical or asynchronous immune and microbiota features precipitating a cascade of events leading to respiratory disease in infants.

In recent years the concept of a “neonatal window of opportunity” of exposure-mediated immune priming has emerged as a potential causal factor underlying chronic immune-mediated diseases (12). This window of opportunity represents a promising target for clinical intervention and disease prevention. Frequent or severe respiratory infections are the leading cause of outpatient visits and hospitalizations in children, and premature infants have up to a 50% risk for recurrent cough and rehospitalization in the first year, most frequently associated with viral infections (35-37). Understanding the earliest immune- and microbial-related events in the context of host development, especially those brought on by poorly timed viral or bacterial exposures during infancy, is essential to interrupting a pathologic program that can lead to chronic respiratory morbidity (38-40). Previous reports have modeled the impact of age on microbiota and immunity, and some features of each system are able to predict respiratory outcome (2, 5, 25, 37). These studies do not necessarily consider when during development features of these systems emerge, or the associated risk or protection conferred by such timing.

Exposures that accelerate or delay the normal maturation of T cells and microbiota during infancy, such as in utero infection promoting the early occurrence of Tphe5, may disrupt this age-specific balance that has served human evolution so well. Microbiota and T cell developmental indices defined in our study reveal that maturity at term and rate of maturation over the first year are indicators of respiratory risk in the first year of life (PRD). Specifically, precocious immune development in conjunction with an immature microbiota at term corresponds to substantially elevated risk of PRD, while either one of these factors alone has an attenuated effect. This indicates that mistimed or discordant maturation between the microbiome and immune systems in newborns establishes a novel, potentially modifiable, pathway to respiratory disease.

T cells, even those derived during gestation, survive into adulthood and therefore have the capacity to remember, or be imprinted by, early exposures (41, 42). Indeed, subjects who deviated from the typical immune trajectory by entering early into the Treg-enriched state type, Tphe5, were more likely to have been exposed to immune- and microbiota-modulating stimuli (chorioamnionitis and antenatal antibiotics). Previous nonhuman primate studies show that intrauterine inoculation with LPS or infection causes expansion of dysregulated FOXP3+ CD4+ T cells (43). Our previous and other studies examining immune responses in human infants exposed to chorioamnionitis or funisitis show an altered placental microbiome as well as sustained inflammatory transcription factor profile, including defects in FOXP3+ T cell function (44, 45). Infants exposed to chorioamnionitis also appear to be at risk for respiratory morbidity (46). Our IST grouping further shows Tregs and effectors are expanded within the same Tphe5 IST, suggesting a state of dysregulation rather than immune suppression. In support of this interpretation, the CCR7-FOXP3+ CD4+ subpopulation associated with Tphe1 arises in inflammatory states and can contribute to immune dysregulation in CCR7 null mice (47). We did not directly test the function of chorioamnionitis-or antibiotic-exposed Tregs, but it is plausible that in our cohort, an aberrant Treg/effector state type shapes both immediate immune responses to infection as well as the ability for protective microbiota state types to take hold. This hypothesis is supported by the observation that Treg-enriched Tphe5 immune state type did not directly predict respiratory morbidity, but did predict a respiratory morbidity-associated pattern of microbiota colonization. It is therefore reasonable to speculate that the impact of antenatal inflammation on later respiratory morbidity is mediated through an aberrant developmental immune-microbiota axis. Interestingly, Tphe5 also predicted CMV infection and the CMV-associated Tphe6 “exhausted” T cell phenotype, which raises the possibility of a complex, bidirectional immune, bacterial and viral cascade that determines infant health outcomes.

Temporal relationships revealed by our longitudinal systems-based approach offer insight into whether or not the risk-associated state types, either immune-or microbiota-related, are modifiable. For example, *Alloiococcus* is substantially diminished during acute respiratory illness reveals a previously undescribed sequence initiated by perinatal inflammation, followed by aberrant T cell phenotype at birth (Tphe5), subsequent airway colonization (low *Alloiococcoccus*-dominated CST8), and ultimately susceptibility to respiratory infection throughout the first year of life. While the antenatal exposures are not easily prevented, there may be some benefit to targeting probiotic treatments, perhaps with *Alloiococcus* species, to interrupting this cascade in chorioamnionitis-exposed mothers and neonates. In fact, probiotics have been successful in several studies in the prevention of illnesses including reductions in pediatric upper respiratory tract infections (48-50). On the other hand, a more conservative, informed approach to immune-or microbiota-based therapy in infants may be called for by a recent report showing that probiotic treatment in healthy newborns had only a transient effect on stool microbiota, and was associated with an increased risk of enteric and respiratory infections (51). Indeed, evidence from our study indicates that the lack of benefit found in probiotic preventive studies may be due to forcing the introduction of microbes into a niche that is appropriate during infant development in general, but is excluded on the basis of an individual infant’s immunological state. Our results underscore the need to assess microbiota and immune systems within the context of one another and the infant’s physiologic development before considering interventions that could disrupt an established, age-appropriate balance.

Our results show that substantial changes in the immune system occur between NICU discharge and 12 months, and in coordination with the gut and respiratory microbiota. More intensive interim sampling, while difficult to perform in infants, is likely to reveal additional factors impacting an individual’s immune trajectory. However, the detailed characterization of relationships between T cell populations and microbiota, and the demonstrated associations between the development of these systems and infant health, represent novel insights into the clinical relevance of microbiome-immune co-development and will inform causal models and mechanistic hypotheses that can be used to develop innovative interventions and guide treatment decisions by furthering understanding of the microbiome-immune axis and the neonatal window of opportunity.

## Materials and Methods

### Study Design

All study procedures were approved by the University of Rochester Medical Center (URMC) Internal Review Board (IRB) (Protocol # RPRC00045470 & 37933) and all subjects’ caregivers provided informed consent. The infants included in the study were enrolled within 7 days of life for the University of Rochester Respiratory Pathogens Research Center PRISM and were cared for in the URMC Golisano Children’s Hospital. Clinical data including nutrition, respiratory support, respiratory symptoms, medications, comorbidities, were entered into REDCap (52, 53), then integrated with laboratory results using the URMC Bio Lab Informatics Server, a web-based data management system using the open source LabKey Server (54). Blood was collected at birth, time of NICU discharge or 36-42 weeks PMA (whichever occurred first), and at 12 months of life. We collected 2729 gut (842 from NICU and 1887 post-discharge), and 2210 nasal (619 from NICU and 1591 post-discharge) usable microbiota samples longitudinally from 139 pre-term and 98 full-term infants and worked with the most extensive subset of these possible depending on the analysis in question (Supplementary Table 2). From the PRISM study cohort, fecal (rectal) and nasal material was collected from pre-term infants (23 to 37 weeks gestational age at birth (GA)) weekly from the first week of life until hospital discharge, and then monthly through one year of gestationally corrected age. Rectal and nasal samples were collected from full-term infants at enrollment and monthly through one year. Additionally, rectal and nasal samples were collected from all infants whenever they exhibited symptoms of acute respiratory illness after discharge from the hospital. Symptoms of acute respiratory illness prompting sample collection were summarized by the primary caregiver using a symptom COAST (Childhood Origins of Asthma) score sheet (35). Parents were instructed to notify the study team if the infant had symptom score of three or greater. Among subjects who completed study procedures through 12 months, 52 PT subjects (43%) and 17 FT subjects (21%) met the criteria for PRD. All blood samples generating usable data were included in all analyses.

Sufficient blood to perform T cell phenotyping by flow cytometry at three pre-defined timepoints was collected from 55% of subjects at birth, 61% of subjects at NICU discharge, and 38% at 12 months. For training the PMA predictions models (described below), all microbiota samples were used. For all other analyses, microbiota samples from subjects that did not have any usable data from blood were excluded. Two staining panels, covering i) intracellular cytokine production (ICS) and ii) T cell surface phenotyping (Tphe) were designed (Supplementary Table 3). Complete immunophenotyping for all three timepoints was performed on 25% of subjects, and 63% of subjects had complete immunophenotyping for at least one timepoint.

### Flow Cytometry Methods

Sample collection, isolation, storage, thawing, stimulation and staining for flow cytometry was performed as detailed previously (55). In short, cord blood and peripheral blood mononuclear cells were isolated via Ficoll centrifugation, cryopreserved and stored in liquid nitrogen, and rapidly thawed and washed with pre-warmed RPMI-1640 (10% FBS and 1x L-glutamine); thawing was done in ‘subject-balanced’ batches (equal mix of pre and full-term subjects, each with three time points) and an aliquot of each freshly thawed sample was plated and stained with a T-cell phenotyping (‘Tphe’) panel with the remainder of the sample rested overnight in an incubator, plated and stimulated with *Staphylococcus aureus*, Enterotoxin Type B (SEB), and stained with a T-cell functional panel (‘ICS’). Panel compositions are as shown in Supplementary Table 3.

Samples were acquired on a BD LSRII (core facility instrument QC-ed daily with BD CS&T beads); PMT voltages normalized per run to pre-determined/optimized ‘Peak-6’ (Spherotech) median fluorescence values. R-based packages and scripts were used for all post-acquisition processing and analysis. Reading of raw .fcs files, compensation, transformation, and subsetting/writing of .fcs files was performed using flowCore (56). To minimize inter-run variation associated with the Tphe panel, the flowStats (57) warpSet function was used to normalize arcsinh transformed channel data using a standard healthy donor adult PBMC control across batches as a reference. For analysis with the clustering algorithm FlowSOM, an iterative approach was used for both panels to first cluster on live, intact, lymphoid-sized CD4+ and CD8+ T-cell subsets (in the case of the ICS panel, including activated, CD69+ subsets); those subsets were then re-clustered to capture rare populations and optimally resolve phenotypic heterogeneity and associated function. Over-clustering followed by expert-guided merging was favored when defining the final number of T cell populations. FlowSOM clustering results used in downstream analysis were represented as proportion of the respective T-cell subset, per sample. All scripts, including Tphe arcsinh cofactors, warpSet and FlowSOM parameters, and final clustering counts are available in Supplementary R-Code.

### Microbiota Identification

Microbiota sample collection and storage techniques, genomic DNA extraction and background control methods were as previously published (58). Raw data from the Illumina MiSeq was first converted into FASTQ format 2□×□312 paired-end sequence files using the bcl2fastq program (v1.8.4) provided by Illumina. Format conversion was performed without de-multiplexing, and the EAMMS algorithm was disabled. All other settings were default. Samples were multiplexed using a configuration described previously (59). The *extract_barcodes*.py script from QIIME (v1.9.1) (60) was used to split read and barcode sequences into separate files suitable for import into QIIME 2 (v2018.11) (61) which was used to perform all subsequent read processing and characterization of sample composition. Reads were demultiplexed requiring exact barcode matches, and 16S primers were removed allowing 20% mismatches and requiring a matching window of at least 18 bases. Cleaning, joining, and denoising were performed using DADA2 (62): reads were truncated (forward reads to 260 bps and reverse reads to 240 bps for rectal V3-V4 samples and forward reads to 275 bps and reverse reads to 260 bps for nasal V1-V3 samples), error profiles were learned with a sample of one million reads per sequencing run, and a maximum expected error of two was allowed. Taxonomic classification was performed with naïve Bayesian classifiers trained on target-region specific subsets of the August, 2013 release of GreenGenes (63). Sequence variants that failed to classify to the phylum level or deeper were discarded. Sequencing variants observed fewer than ten times total, or in only one sample, were discarded. Rectal samples with fewer than 2250 reads and nasal samples with fewer than 1200 reads were discarded. Phylogenetic trees were constructed for each body site using MAFFT (64) for sequence alignment and FastTree (65) for tree construction. For the purposes of β-diversity analysis, rectal and nasal samples were rarefied to depths of 2250 and 1200 reads, respectively, and the Unweighted Unifrac (66) metric was applied.

### Cytomegalovirus Detection

Subjects were screened for prior prior or current cytomegalovirus (CMV) exposure, using plasma serology tested at the University of Rochester Clinical Virology Laboratories. Active infection was assessed by Real-Time PCR according to a previously published protocol using a double-primer assay targeting the UL55-UL123-exon 4 regions (67).

### Statistical analyses

#### Multivariate ANOVA

We used a sequence of multivariate ANOVA (MANOVA) models to estimate the amount of variance one set of variables could explain in another. We modeled T cell population relative abundances, gut, and nasal species-level relative abundances pairwise each as predictor and response matrices. DOL and PMA served only as predictors. Each pair of variables types was joined, with missing samples deleted casewise. T cell populations and microbiome taxa with a variance of less than .0001 were removed in each comparison. The remaining variables were renormalized to sum to one, and transformed using the isometric log ratio, then modeled using a multivariate linear model with a matrix response. 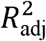 was calculated as 1-MSE_full_ /MSE_reduced_ where the mean squared error (MSE) was the total sum of squared residuals in the response matrix, divided by the residual degrees of freedom, thus approximately unbiased for the residual variance. The PMA-adjusted model used PMA, and the set variables of interest as a predictor in the full model, retaining only PMA in the reduced model. Wilks’ lambda was used to test for association between response and predictor variables.

#### T cell PMA- and GA-associated trajectories

For each T cell subpopulation, a linear regression was fit of GA and PMA on the arcsin-sqrt transformed relative abundance of that population using a continuous and piecewise linear model with a single knot at 37 weeks PMA and interaction with GA (Fig. 2C-2D and Supplementary Fig. 1). In symbols, we fit the model 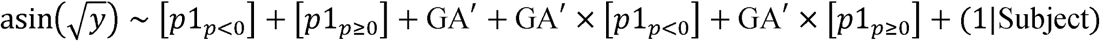, where *y* is the relative abundance of T cell population (relative to other populations that share same CD4 vs CD8 status and Tphe vs ICS), *p* = PMA - 37, GA^’^ = GA - 37 are the PMA and GA of a sample, shifted so that term-equivalent samples and gestational ages are zero, and (1|Subject) is a random intercept for each subject. The intercept of this model represents the abundance in subjects of 37 weeks GA at birth, and the remaining terms are identified by interpolation and extrapolation of the time points actually sampled in an individual. Figure 2D plots the GA^r^ term and its 95% CI for T cell populations with significant (Bonferroni-adjusted p<.05, 80 tests) GA^r^ effects. Non-monotone populations were determined by testing three contrasts i) the *p*1_*p*<O_] terms, ii) the *p*1_*p*≥O_] terms, and iii) the difference between them for joint statistical significance (Bonferroni-adjusted p<.05, 80 tests).

*CST and IST Assembly*. Microbial community state types (CSTs) were defined for each body site by fitting Dirichlet multinomial mixture (DMM) models (68) using the R package DirichletMultinomial (v1.22.0) (69, 70), R version 3.5.0. Sample composition was represented using normalized counts of the most specific operational taxonomic units (OTUs) present in at least 5% of the samples from a given body site. Normalization was performed on a per sample basis by taking the relative abundance of each OTU and multiplying by 2250 for rectal samples and 1200 for nasal samples. Resulting non-integer counts were rounded down. For each body site, the DMM model was fit with one through twenty Dirichlet components and the optimal number of components was selected by minimizing the Laplace approximation of the negative-log model evidence. In this model, CSTs are synonymous with Dirichlet components, and each sample was assigned to the CST from which it had the highest posterior probability of be derived. This procedure was repeated with the immunological data in order to define immune state types (ISTs), using relative abundances of FlowSOM defined T cell populations in the place of OTUs. Relative abundances were computed within assays (TPHE and ICS) and major populations (CD4 and CD8) separately, and converted to counts by multiplying by 50,000 and rounding down. CD4 and CD8 counts were combined to fit the DMM for each assay.

#### Microbiota-T cell Associations

Associations between microbiome development and the immune system were modeled using microbiome CST occurrence patterns as outcome variables and iterating through the relative abundances of each FlowSOM T cell population or observed IST at each time point as predictors. In symbols, we used the model

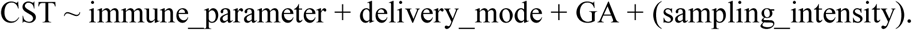

For each CST, each of these immunological parameters (T cell population relative abundances and IST, hereafter referred to as the immunological variables of interest [VOIs]) at each of the three time points when the immune system was sampled (birth, discharge, and one year) was assessed independently. CST occurrence patterns were related to immunological VOIs by testing three types of associations between every CST-VOI combination at the level of individual subjects, while controlling for mode of delivery (MOD), gestational age at birth (GA) and, in model (i) only (see below), the number of microbiome samples (sampling_intensity) that were collected from an individual. These models differed in the aspect of CST occurrence that was modeled as the outcome. Model (i) tests associations between the VOI and whether or not a CST occurs at all in an individual; (ii) tests associations between the VOI and how persistent a CST is in an individual; and (iii) tests associations between the VOI and the days to first occurrence of a CST in an individual. Model (i) was tested using logistic regression with VOI, MOD, GA and the number of microbiome observations from a given individual as the sampling intensity. The outcome indicated whether or not a given CST was ever observed in the individual. We tested the VOI association by dropping that term and calculating a likelihood ratio test. Model (ii) was tested using a quasi-Poisson regression model with MOD, GA, and the VOI as covariates, and total number of days the subject was assigned to *any* CST as an offset. The number of days a subject was assigned to a given CST was the outcome and was calculated by summing the interval lengths between CST change points. Intervals were calculated from midpoint to midpoint on the sampled days of life. At birth, subjects were placed in the first observed CST if the first sample occurred within 14 days of life, otherwise the first interval was excluded. Subjects were assumed to remain in their final observed CST for an interval equal to half the interval length between the penultimate and ultimate sample. Significance of the VOI was assessed as in model (i). Model (iii) was tested using interval censored, accelerated log logistic failure time models (R package icenReg v2.0.9) (71) with MOD, GA, and the VOI as covariates and the interval preceding the first observation of a given CST as the outcome. For gCST 1 and nCST 1, which on average were the earliest CSTs, we modeled the interval preceding the first observation of a CST other than NAS 1 or REC 1. For each CST, only subjects that were ever observed in that CST at some point were included. Significance was assessed based on Wald test p-values of the terms in the fitted full models.

For models (i)-(iii), subjects with fewer than one sample taken per 30 NICU-days or fewer than six samples post discharge were excluded. We filtered immune VOI with fewer than ten observations, and CSTs present in fewer than 10% of the remaining observations. Numerical covariates were converted into z-scores, except GA which we modeled as (GA - 37)/37. Within each model (i)-(iii), multiple testing across all CSTs and VOIs was corrected for using the Benjamini-Hochberg method at 10% FDR.

#### Tphe5, Alloiococcus abundance, and acute illness associations

Using only post-discharge nasal samples, the abundance of *Alloiococcus* represented as read counts was modeled as a function of day of life, GA, MOD, and the occurrence of Tphe5 at birth or discharge using a generalized estimating equation fit with the geeglm function in R (72). Subject was used as the clustering variable, an exchangeable working correlation structure was specified, total reads per sample was used as an offset, and the family was Poisson with a log link function. This model was repeated with the addition of acute illness as a covariate. The probability of a sample coming from an illness or healthy surveillance visit was modeled using mixed effects logistic regression fit with the glmer function (73), using *Alloiococcus* relative abundance, DOL, GA, and MOD as covariates, with Subject as a random effect. This model was repeated with the addition of Tphe5 at birth or discharge as a covariate.

#### Prediction of PMA

Two separate elastic net regression models (74) were trained to predict (75) the log2-transformed PMA with a) T cell immunological features and b) microbial abundance. In (a) the four feature sets were CD4 ICS, CD8 ICS, CD4 Tphe and CD8 Tphe populations, while in (b) the two feature sets consisted of nasal and rectal species-level relative abundances from samples collected prior to DOL 450, filtered to remove taxa present in fewer than 3% of samples. A total of 433 samples from 185 subjects and 80 features were included in (a). Model (b) was trained on 3032 samples from 237 subjects and 218 features. Some samples had incomplete feature sets, e.g., if only the ICS panel was run then both the CD4 and CD8 Tphe sets were missing, or if only the nasal microbiome was sampled and the rectal abundances were missing. We treated this as a missing data problem, and imputed the values with their mean values among non-missing cases. Imputation was chained onto the elasticnet model (occurred only using the training data, in each fold) for the purposes of tuning and validation. Within each feature set, we used the relative proportions, transformed into z-scores.

#### Cross validation for tuning and prediction

We tuned the model and estimated its performance using cross-validation by holding out a subject’s entire longitudinal record. We tuned the elastic net alpha in [0, 1] and lambda in [.001, .5] parameters by randomly selecting 50 combinations of (alpha, lambda) and evaluating the test mean-square error (MSE) via 5-fold cross-validation. After finding a minimizing pair of (alpha, lambda), the model was refit with 10-fold cross-validation. For each subject i, this provides two sequences of fitted values, representing the log2-transformed PMA prediction. For instance, for the microbiome, we have

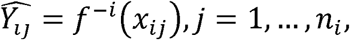

where *x*_*ij*_ represent microbial feature vectors, *n*_*i*_ indexes the number of longitudinal samples for subject *i*, and *f*^-*i*^ represent the elastic net model trained excluding subject i. For the T cell immunome, the analogous model is fit. The back-transformed values 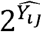 were used to calculate each model’s *R*^2^.

#### Immunological and microbial developmental indices

The longitudinal sequence of cross-validated fitted values 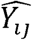 were compared to the true PMA for each subject using a linear mixed model. We fit the model

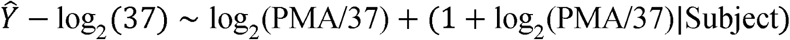

thus 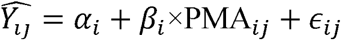 and calculated the best linear unbiased predictor of each subject’s 37-week intercept *α*_*i*_, slope *β*_*i*_ and their conditional standard errors se(*α*_*i*_), se(*β*_*i*_). These are transformed into a quantity similar to a z-score by subtracting the median of *α*_*i*_, *β*_*i*_ over subjects i, and dividing by its conditional standard error se(*α*_*i*_) or se(*β*_*i*_).

#### Prediction of PRD

We used random forest classification models to predict PRD using two feature sets: clinical and developmental index. The clinical features were race, maternal education, the baby’s sex, gestational age, weight and season at birth, and oxygen supplementation integrated over the first 14 days of life. The developmental index features were the z-scores of the microbiome and T-immune slopes and intercepts. The random forest hyperparameters mtry, ntree and nodesize were tuned separately for each feature set with random search using 5-fold cross-validation. After the optimal parameters were found for each feature set, a second round of 20-fold cross validation was used to evaluate the area under the ROC curve (AUC). The fitted values from the random forest regression were calculated using the function generatePartialDependenceData.

## Supporting information

Combined supplemental tables and figures

## Acknowledgments

Authors would like to acknowledge the University of Rochester Pediatrics Translational Biospecimen Laboratory, Genomics Research Core, Flow Cytometry Core, and the Human Immunology Center. We also thank Richard Simon and Kimberly Baldo (The Harley School), for their assistance with figure preparation and graphic design. We thank the nurses and staff at the Golisano Children’s Hospital NICU and URMC Strong Beginnings Maternity Services, and most of all our families who generously consent to research studies. Funding was provided by NIH NIAID HHSN272201200005C (Respiratory Pathogens Research Center), NIH NIAID 1K08AI108870-01A1 (CD8 T Cell Dysregulation in Premature Infants), NIH NHLBI U01 HL101813-01 (Prematurity and Respiratory Outcomes Program), NIH NCATS UL1 TR000042 (Clinical and Translational Science Institute).

## Author Contributions

Conception, execution, interpretation and manuscript preparation: KMS, AM, AG, NL, GSP, MC, AD

Project PI’s, University of Rochester: GSP, MC, SG, AF, DJT

Acquisition and analysis of experimental data: NL, AG, ALG, HK, JC, KMS

Clinical and sample data collection, study coordination and recruitment: EC, HH, GSP, MC, KMS

Biostatistical analysis: AM, AG

Clinical and laboratory data integration and management: JHW, SB

## Data and Materials Availability

Annotated datasets for 16S sequencing and flow cytometry results can be found in dbGaP, accession number phs001347. Code supporting this paper is available at https://github.com/amcdavid/CoordinatedTCellsMicrobiota.

