## Supplementary material for "Aberrant newborn T cell and microbiota developmental trajectories predict respiratory compromise during infancy": Combined supplemental tables and figures

| **TPHE** | subjects | samples |
|---|---|---|
| 3 timepoints | 78 | 234 |
| 2 timepoints | 82 | 164 |
| 1 timepoint | 16 | 16 |
| total | **176** | **414** |

| **TPHE** | samples | pre-term | full-term |
|---|---|---|---|
| birth | 147 | 65 | 82 |
| discharge | 163 | 87 | 76 |
| 12-month | 104 | 58 | 46 |
| total | **414** | *210* | *204* |

| **ICS** | subjects | samples |
|---|---|---|
| 3 timepoints | 69 | 207 |
| 2 timepoints | 89 | 178 |
| 1 timepoint | 19 | 19 |
| total | **177** | **404** |

| **ICS** | samples | pre-term | full-term |
|---|---|---|---|
| birth | 147 | 64 | 83 |
| discharge | 155 | 84 | 71 |
| 12-month | 102 | 55 | 47 |
| total | **404** | *203* | *201* |

| **TPHE_ICS** | subjects | samples |
|---|---|---|
| 3 timepoints | 67 | 201 |
| 2 timepoints | 83 | 166 |
| 1 timepoint | 18 | 18 |
| total | **168** | **385** |

| **TPHE_ICS** | samples | pre-term | full-term |
|---|---|---|---|
| birth | 141 | 63 | 78 |
| discharge | 150 | 81 | 69 |
| 12-month | 94 | 51 | 43 |
| total | **385** | *195* | *190* |

**Supplementary Table 1.** Subject numbers for immunophenotyping

| Figure Description | Site | Samples | Subjects |
|---|---|---|---|
| **Microbiome CGA & CST PCoA plots** | Both NAS | 1748 | 149 |
| | Both REC | 1899 | 143 |
| **Microbiome CST Occurrence Over CGA** | NAS | 1748 | 149 |
| | REC | 1899 | 143 |
| **Microbiome Composition Heatmaps** | NAS | 1748 | 149 |
| | REC | 1899 | 143 |
| **Immuno IST Composition Heatmaps** | TPHE | 414 | 176 |
| | ICS | 404 | 177 |
| **Immuno IST Occurrence Over CGA** | TPHE | 414 | 176 |
| | ICS | 404 | 177 |
| **Immuno IST Avg. Occurrence GAB/CGA** | TPHE | 414 | 176 |
| | ICS | 404 | 177 |
| **NAS 8 Occurrence vs TPHE ISTs** | Birth | 68 | 68 |
| | Discharge | 95 | 95 |
| **CST-Immuno Association Networks** | NAS | 1589 | 109 |
| | REC | 1697 | 117 |

**Supplementary Table 2.** Subject numbers microbiome and combined analyses

| Cytometer: BDLSRII (URMC FlowCore - Animal) | | | | | | | | | |
|---|---|---|---|---|---|---|---|---|---|
| Tphe Functional Panel (RPRC 12-0012) | | | | | | | | | |
| **Laser** | **Long Pass** | **Band Pass** | **PMT** | **Detector** | **Marker** | **Color** | **Clone** | **Company** | **Catalog #** |
| 488 | 505 | 515/20 | BB | B515 | **CD122** | BB 515 | Mik-β | BD Biosciences | 564688 |
| 488 | 685 | 710/50 | BA | B710 | **Perforin** | PerCP-Cy5.5 | dG9 | Biolegend | 308114 |
| 407 | | 450/50 | VH | V450 | **Granzyme B** | BV 421 | GB11 | BD Biosciences | 563389 |
| 407 | 535 | 550/40 | VG | V550 | **Live/Dead** | Aqua | | Life Technologies | L34957 |
| 407 | 570 | 585/42 | VE | V585 | **CD3** | BV 570 | UCHT1 | Biolegend | 300436 |
| 407 | 595 | 605/40 | VD | V605 | **CD31** | BV 605 | WM59 | BD Biosciences | 562855 |
| 407 | 630 | 660/40 | VC | V660 | **CD127** | BV 650 | HIL-7R-M21 | BD Biosciences | 563225 |
| 407 | 670 | 705/70 | VB | V705 | **CD45RO** | BV 711 | UCHL1 | BD Biosciences | 563722 |
| 407 | 740 | 780/60 | VA | V780 | **CD8a** | BV 785 | RPA-T8 | Biolegend | 301045 |
| 633 | | 660/20 | RC | R660 | **KLRG1** | APC | 13F12F2 | eBioscience | 17-9488-42 |
| 633 | 685 | 710/50 | RB | R710 | **CD185 (CXCR** | APC-R700 | RF8B2 | BD Biosciences | 565191 |
| 633 | 740 | 780/60 | RA | R780 | **CD197 (CCR7** | APC-Cy7 | G043H7 | Biolegend | 353212 |
| 532 | | 575/24 | GE | G575 | **Foxp3** | PE | 236A/E7 | eBioscience | 12-4777-42 |
| 532 | 600 | 610/20 | GD | G610 | **CD4** | PE-TR | S3.5 | Invitrogen | MHCD0417 |
| 532 | 640 | 660/40 | GC | G660 | **CD28** | PE-Cy5 | CD28.2 | BD Biosciences | 561791 |
| 532 | 740 | 780/40 | GA | G780 | **CD57** | PE-Cy7 | TB01 | eBioscience | 25-0577-42 |

| ICS Functional Panel (RPRC 12-0012) | | | | | | | | | |
|---|---|---|---|---|---|---|---|---|---|
| Cytometer: BDLSRII (URMC FlowCore - Animal) | | | | | | | | | |
| **Laser** | **Long Pass** | **Band Pass** | **PMT** | **Detector** | **Marker** | **Color** | **Clone** | **Company** | **Catalog #** |
| 488 | 505 | 515/20 | BB | B515 | **IL-8** | FITC | E8N1 | BioLegend | 511406 |
| 407 | | 450/50 | VH | V450 | **IL-17** | Pacific Blue | BL168 | BioLegend | 512312 |
| 407 | 535 | 550/40 | VG | V550 | **Live/Dead** | Aqua | polyclonal | Life Technologies | L34957 |
| 407 | | | | | **CD14** | BV510 | MϕP9 | BD Biosciences | 563079 |
| 407 | 570 | 585/42 | VE | V585 | **CD8a** | BV570 | RPA-T8 | BioLegend | 301037 |
| 407 | 595 | 605/40 | VD | V605 | **IL-2** | BV605 | MQ1-17H12 | BD Biosciences | 564165 |
| 407 | 630 | 660/40 | VC | V660 | **CD45RA** | BV650 | HI100 | BD Biosciences | 563963 |
| 407 | 670 | 705/70 | VB | V705 | **IL-10** | BV711 | JES3-9D7 | BD Biosciences | 564050 |
| 407 | 740 | 780/60 | VA | V780 | **TNFa** | BV785 | MAb11 | BioLegend | 502948 |
| 633 | | 660/20 | RC | R660 | **IL-6** | APC | MQ2-13A5 | BD Biosciences | 561441 |
| 633 | 685 | 710/50 | RB | R710 | **CD3** | AF700 | UCHT1 | BD Biosciences | 557943 |
| 633 | 740 | 780/60 | RA | R780 | **CD69** | APC-Cy7 | FN50 | BioLegend | 310914 |
| 532 | | 575/24 | GE | G575 | **IL-4** | PE | MP4-25D2 | | |
| 532 | 600 | 610/20 | GD | G610 | **CD107a** | PE-CF594 | H4A3 | BD Biosciences | 562628 |
| 532 | 690 | 710/50 | GB | G710 | **CD4** | PE-Cy5.5 | S3.5 | ThermoFischer | MHCD0418 |
| 532 | 740 | 780/40 | GA | G780 | **IFN-g** | PE-Cy7 | B27 | BD Biosciences | 557643 |

**Supplementary Table 3.** Flow Cytometry Panels

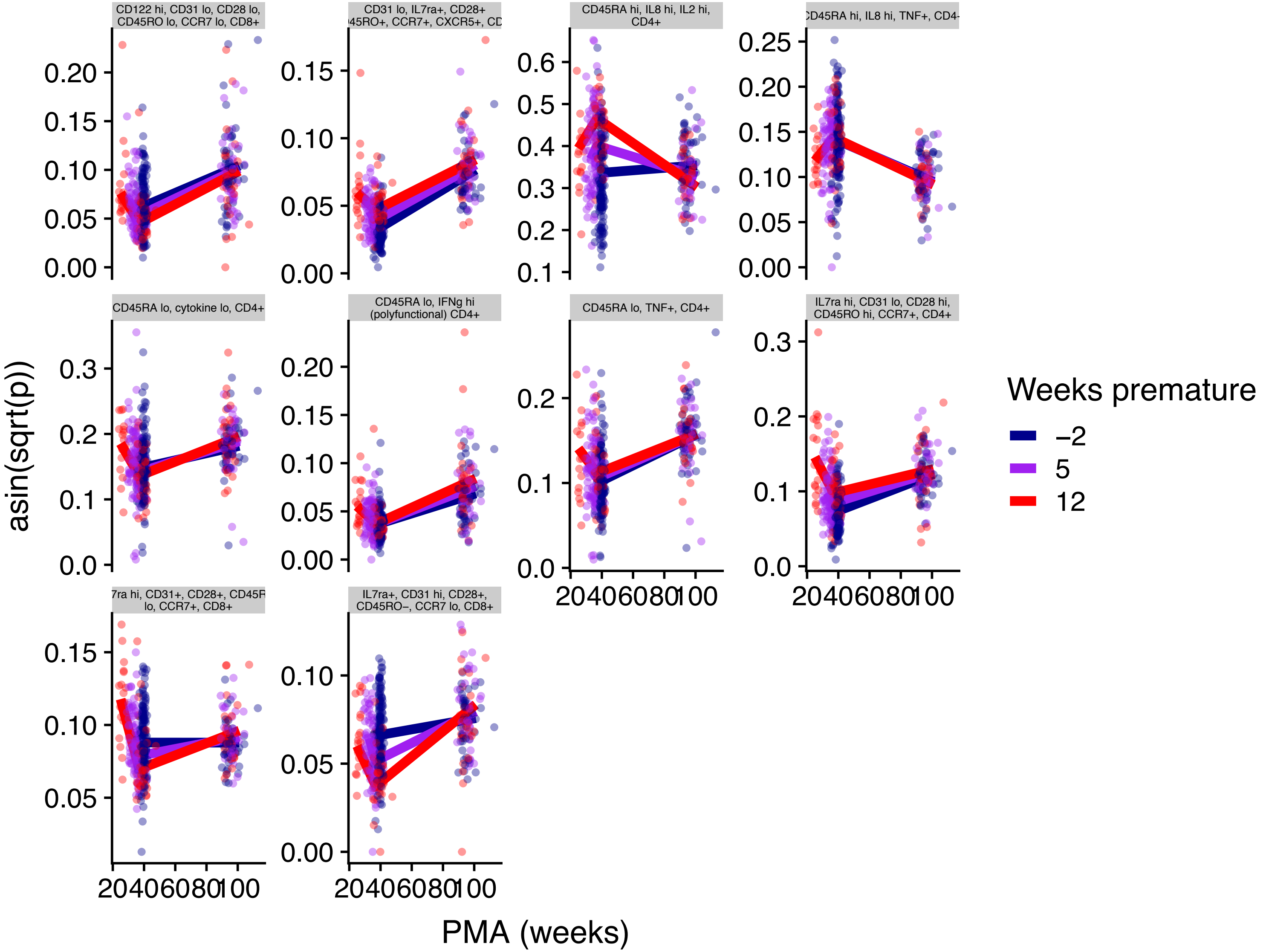


**Supplementary fig 1. Meta clusters with non-monotone trajectories as a function of PMA.**

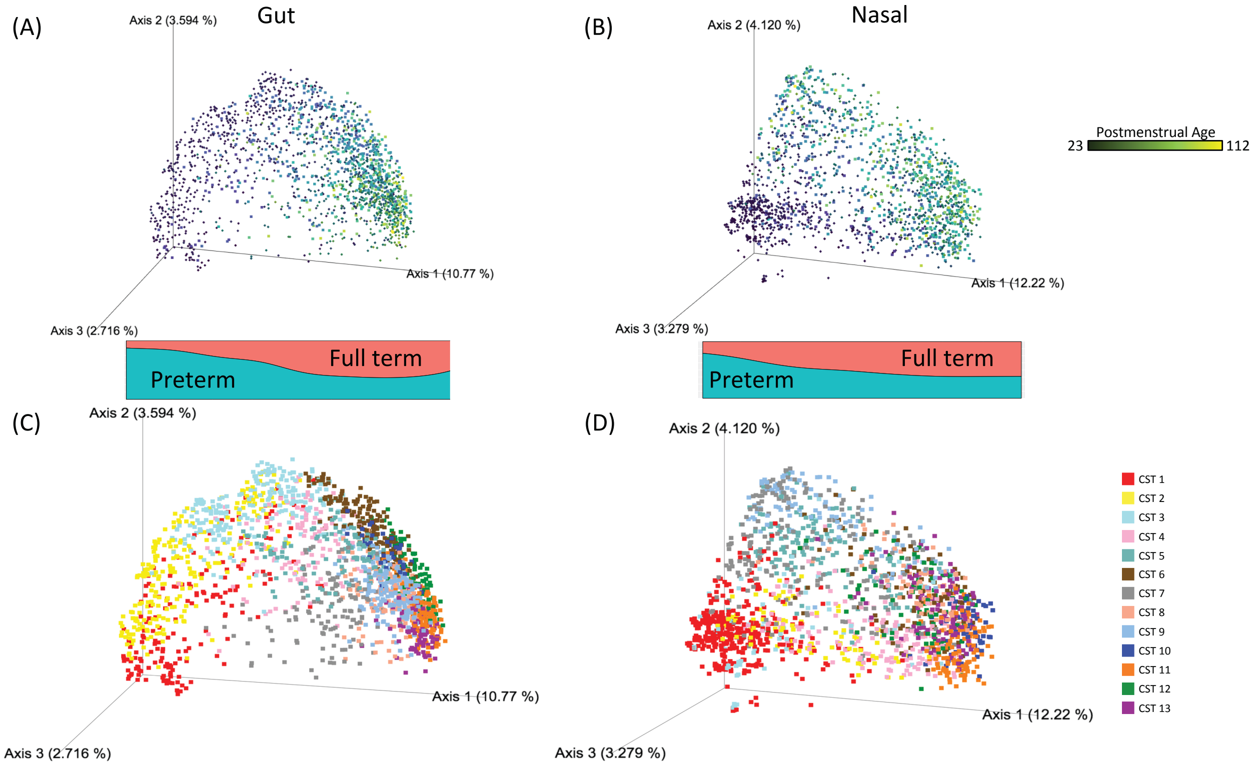


**Supplementary fig 2.** Premature birth influences long-term age-related respiratory and gut microbiota community progression. Microbiota community profiling was performed on rectal (A, C) and nasal (B, D) samples obtained from 159 infants during regular surveillance and acute respiratory illness. (A-D) Principal coordinate analysis (PCOA) plots using Unweighted Unifrac distances summarize overall variation and structure. (A-B) Points were colored by postmenstrual age (PMA) at the time the sample was obtained. Colored bands at the base of PCOA plots show the proportion of samples along each point of axis 1 that are from either preterm (teal) or full term (salmon) subjects. (C and D) Microbiota community state types (CST) were defined for each body site, with samples in the PCOA colored according to the CST they represent. CSTs are ordered according to average PMA of occurrence.

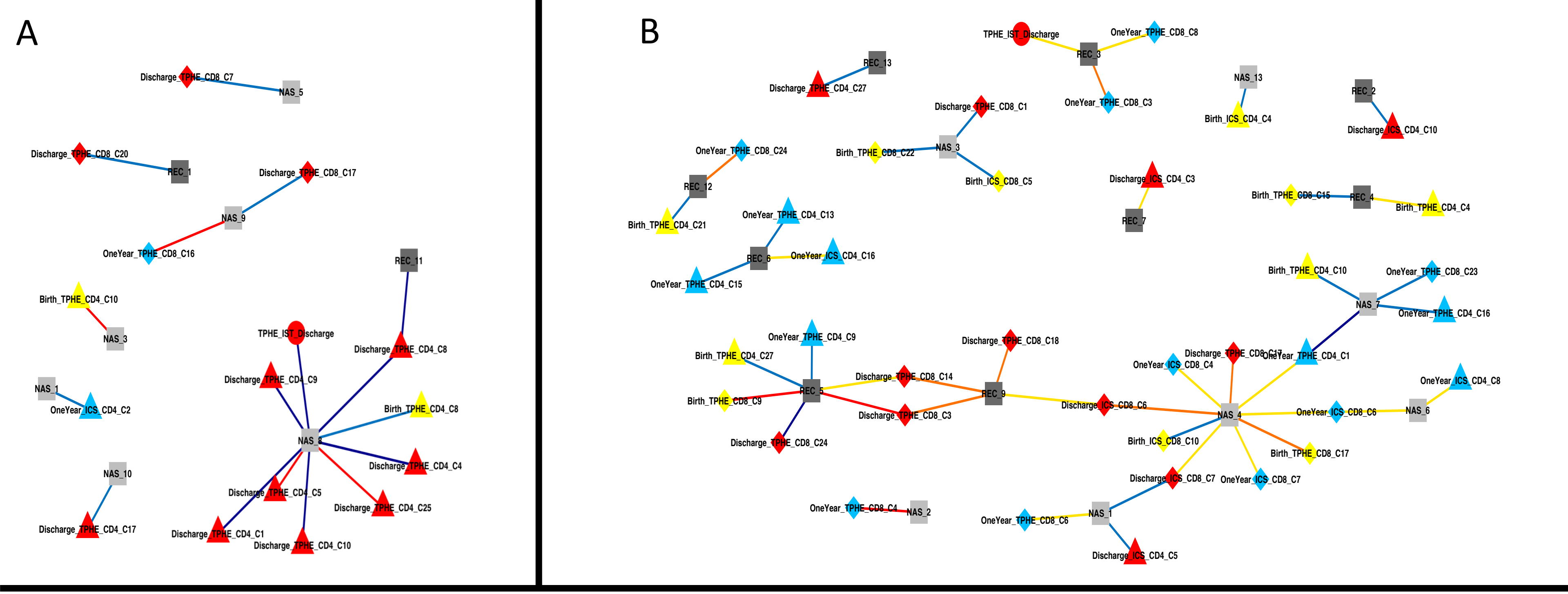


**Supplemental fig 3. Networks of associations between CST occurrence and T cell populations and ISTs.** (A) Logistic regression was used to assess the relationship between T cell populations or ISTs at birth, discharge, or one year and the probability of ever observing a given CST within a subject. (B) Interval censored survival modeling was used to assess the relationship between T cell populations and ISTs at birth, discharge, or one year and the time to occurrence of a given CST within a subject. Mode of delivery and gestational age at birth were included as covariates in both types of models. All significant associations (after multiple test correction) are plotted, with edges between an immunological parameter and a CST indicating a significant relationship. Edges are colored according to the direction of the relationship and the magnitude of its significance. Nodes are colored to indicate time point or body site and are shaped to distinguish between CSTs, ISTs, and individual T cell populations.

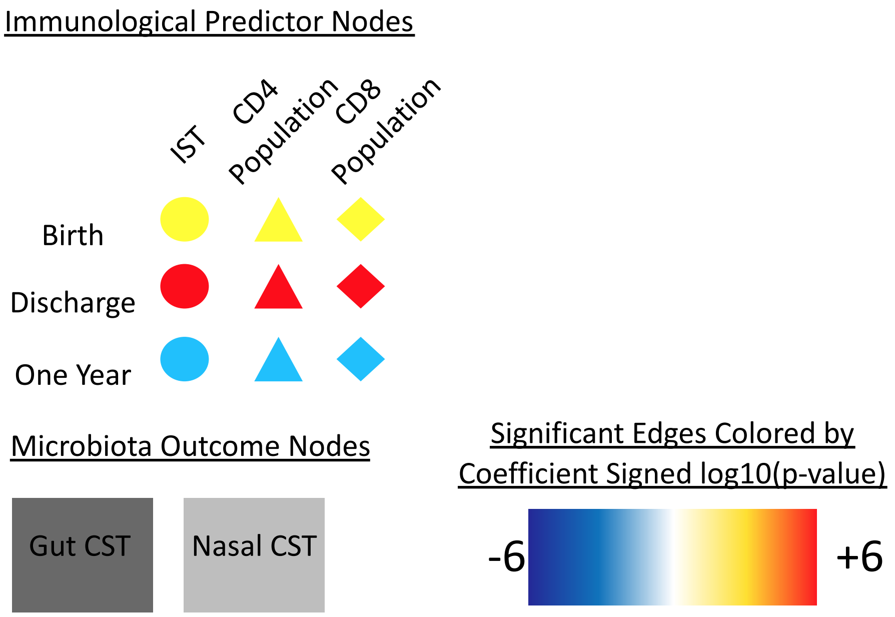

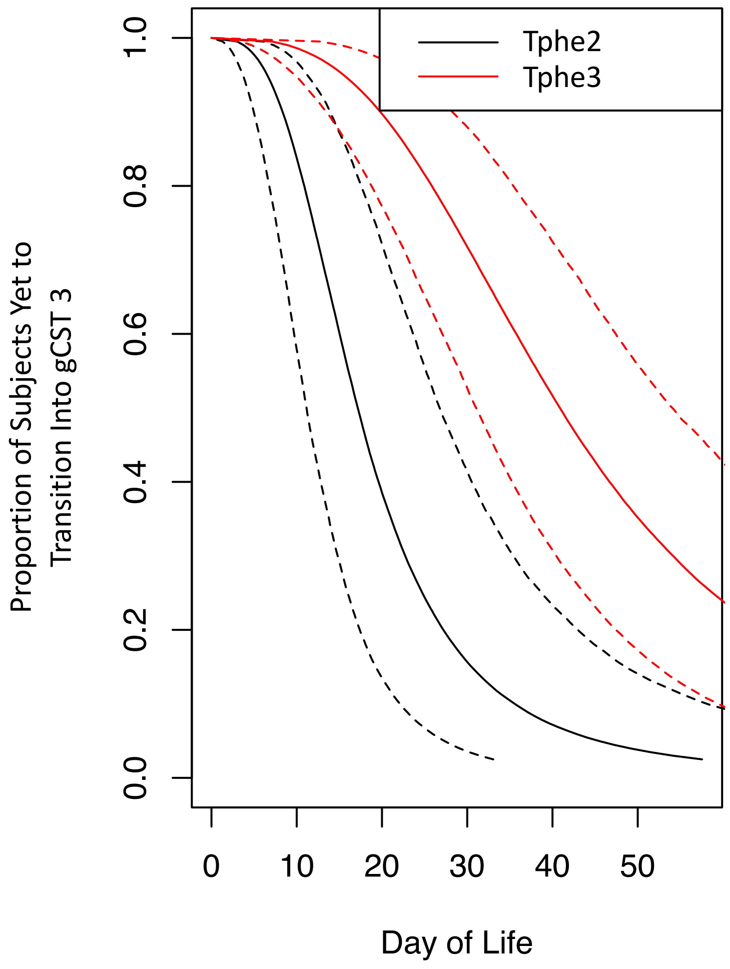


**Supplemental fig 4. Time to transition into gCST 3 based on Tphe IST at discharge.** Survival analysis using an accelerated failure time model was used to assess the time to initially transition into gCST 3 as a function of Tphe IST at discharge, gestational age at birth, and mode of delivery. Fitted mean probabilities of not having transitioned in gCST 3 are shown for infants born at 30 weeks GA by Cesarean section, with 95% confidence intervals. Tphe3 at discharge significantly delayed initial transition into gCST 3.

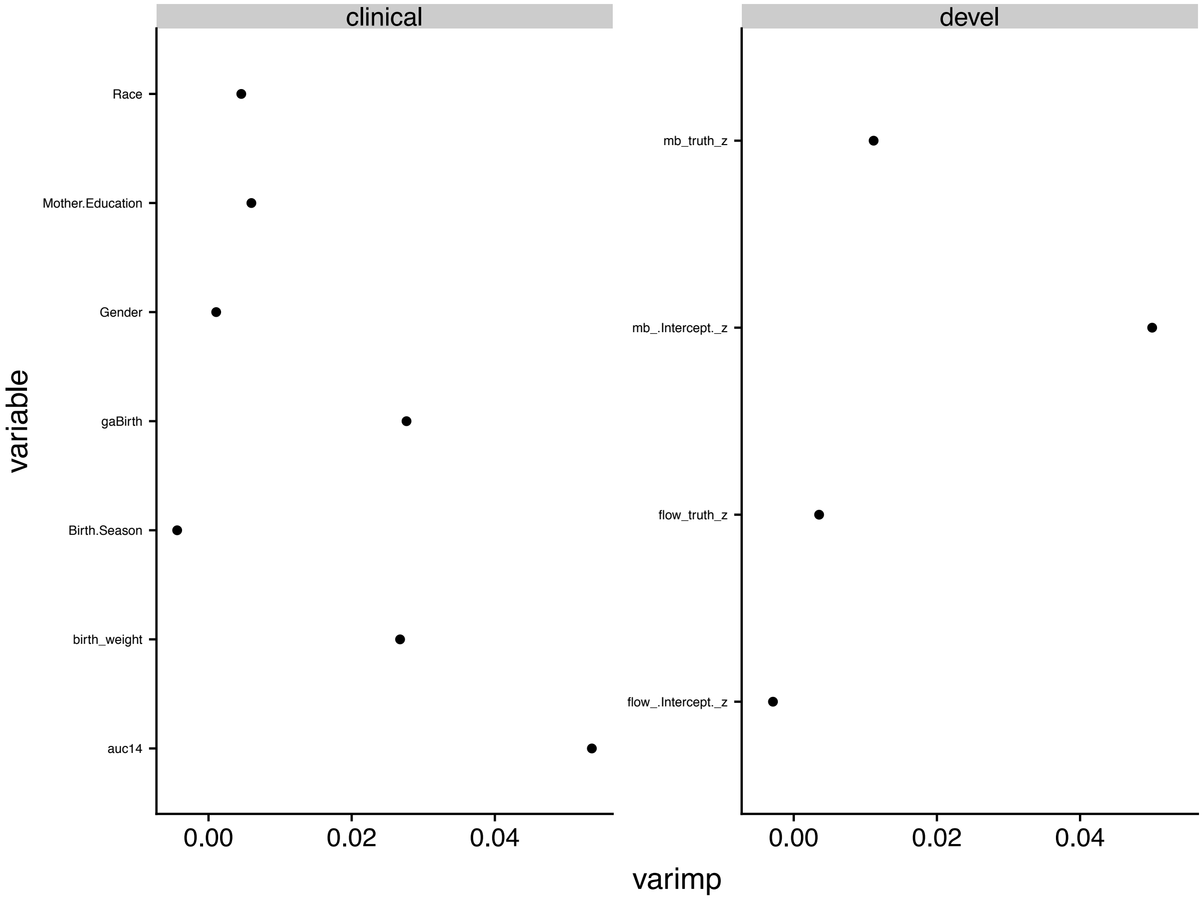


**Supplementary fig 5. Random forest variable importance plots for clinical and developmental index models**. Larger values represent greater decreases in the Gini purity coefficient. Importance was calculated using the default method in R package randomForestSRC version 2.7.0.
